# Disturbed mitochondrial energy production in methylmalonic aciduria is cell-type and variant-dependent

**DOI:** 10.64898/2026.09.23.753738

**Authors:** Miriam A. Güra, Pablo Carbajal Martin, Lisa Tidecks, Caroline T. Glatthard, Julia Lis, Matthew C.S. Denley, Céline Bürer, Seraina Lutz, Anke Schumann, Stefan Kölker, Olivier Devuyst, Florian Traversi, Raphael J. Morscher, D. Sean Froese, Matthias R. Baumgartner

## Abstract

Methylmalonic aciduria (MMA) is caused by deficiency of methylmalonyl-CoA mutase (MMUT), which catalyses the final step of propionate catabolism and fuels the tricarboxylic acid cycle for energy production. Previous studies reported disrupted mitochondrial homeostasis in MMA, including reduced mitochondrial membrane potential and increased oxidative stress, especially in energetically demanding and chronically affected tissues like brain and kidneys. However, how these changes impact mitochondrial energy production remains unclear. Here, we systematically investigated mitochondrial energy production using extracellular flux analysis in cellular models of MMA, including 293T cells, patient-derived fibroblasts, urine-derived epithelial kidney cells and induced pluripotent stem cells (iPSCs) as well as iPSC-derived neurons harbouring either complete loss (knockout, KO) or pathogenic missense variants in MMUT. We found no impact on mitochondrial energy production in MMUT-KO 293T cells and fibroblasts compared to controls. In contrast, fibroblasts and 293T cells expressing the pathogenic MMUT-p.N219Y variant showed decreased energy production. This corresponds to the altered mitochondrial membrane potential in 293T cells, but contrasts with their unchanged mitochondrial abundance. Depletion of glucose, glutamine and pyruvate from the media or provision of each individually as sole fuel source exacerbated the phenotype of MMUT-p.N219Y clones, but did not induce a phenotype in MMUT-KO 293T cells. Finally, we confirmed reduced mitochondrial energy production in patient-derived kidney cells but found no evidence in MMUT-p.N219Y iPSCs and their derived neurons. Overall, our work suggests that the impact of MMUT-deficiency on mitochondrial energy production is cell type- and variant-dependent. Further investigations should clarify molecular mechanisms and their clinical impact.

**Synopsis:** Using extracellular flux analysis across multiple MMUT-deficient cell models, this study shows that impaired mitochondrial energy production occurs with the missense variant p.N219Y but not with complete MMUT loss, and that this effect varies by cell type.

## 1. Introduction

Isolated methylmalonic aciduria (MMA) is an autosomal-recessive inborn error of metabolism caused by deficiency of the enzyme methylmalonyl-CoA mutase (MMUT; MIM #251000) or enzymes involved in the synthesis of its cofactor adenosylcobalamin (AdoCbl)^1,2^. Individuals affected by MMA experience acute symptoms of vomiting, hypotonia, lethargy and life-threatening metabolic decompensation, accompanied by biochemical findings of metabolic acidosis, ketosis and hyperammonemia. Chronic symptoms of MMA are especially found in high-energy consuming organs, manifesting as neurological complications such as stroke-like episodes, seizures, movement disorders and developmental delay^2–4^, as well as impaired kidney function manifesting as chronic kidney disease leading to end stage renal disease^2,5,6^.

MMUT catalyzes the conversion of methylmalonyl-CoA to succinyl-CoA, a step required for the breakdown of branched-chain amino acids, odd-chain fatty acids, cholesterol, and gut microbiome-derived propionic acid, which fuels the tricarboxylic acid (TCA) cycle for mitochondrial energy production. Disruption of this pathway by MMUT-deficiency leads to the accumulation of propionyl-CoA and methylmalonyl-CoA and their potentially toxic derivatives methylmalonic acid, 2-methylcitrate and 3-hydroxypropionic acid. It has been suggested that the accumulation of these intermediate metabolites and/or energy depletion drive disease pathogenicity^3,7–9^, particularly by disturbing mitochondrial processes^10–13^.

Multiple cellular and animal models have been used to investigate how mitochondrial architecture and function are impacted in MMA. Primary skin-derived fibroblasts are widely used for diagnostic purposes and have revealed disturbed anaplerosis of the TCA cycle as a relevant pathomechanism of MMA^9^. Another widely reported characteristic of MMA is mitochondrial dysfunction, based on MMUT being localized in the mitochondrial matrix and its proximity to TCA cycle enzymes^9^. Previous studies showed aberrant morphology (including disorganized cristae and megamitochondria)^6,10,11,14^ and reduced mitochondrial membrane potential^10,12^ in liver tissue from *Mmut*^-/-^ mice^11^, murine^6^ and human kidney tissue^14^, neurons derived from human induced pluripotent stem cells (iPSCs)^12^ as well as urine-derived epithelial kidney cells^10^. Impaired mitophagy^10^, increased reactive oxygen species^10^ and oxidative stress^10,11,13^ were found in the same epithelial kidney cells. Furthermore, altered mitochondrial DNA (mtDNA) to nuclear DNA (nDNA) ratios were detected in these cells^10^ as well as hepatocytes from human and murine liver extracts^15^ and reduced activity of isolated complexes of oxidative phosphorylation (OXPHOS) was measured in murine hepatic extracts^11^ and human tissue biopsies^16^. Finally, decreased oxygen consumption has been reported in urine-derived epithelial kidney cells^10,17^ and in a pilot study with lymphocytes from two individuals affected by MMA^18^. However, despite these findings, the consequences of MMUT-deficiency across tissues and cell-types on mitochondrial energy production have not yet been systematically evaluated.

Here we used extracellular flux analysis, a method widely used to assess mitochondrial function^19^, to measure energy production in MMUT-deficient cells. We examined cells with complete absence of MMUT as well as loss of enzyme function due to pathogenic missense variants across different energy sources and cell-types with different energetic needs. Overall, we found that mitochondrial energy production is highly influenced by the provided fuel source, the cell-type investigated and the underlying disease-causing variant, suggesting a nuanced situation where the impact of MMUT-deficiency on mitochondrial energy production also depends on other factors.

## 2. Material and Methods

### 2.1. Ethics statement

Approval to use patient fibroblasts was granted by the Cantonal Ethics Commission of Zurich (KEK-ZH-Nr. 2014-0211, amendment PB_2020-00053). Urine-derived epithelial kidney cells were provided as described in^17,20^.

### 2.2. Cell culture

293T cells (ATCC, CRL-3216), fibroblasts^9,21^ (control line for iPSC generation: ATCC, CRL-2522) and urine-derived epithelial kidney cells^20^ were cultured in Dulbecco’s modified eagle medium (DMEM; Gibco, 31966-047) supplemented with 10% fetal bovine serum (FBS; Gibco, 102070-106) and 1% Pen/Strep (100X; Gibco, 10378016) at 37 °C and 5% CO_2_ in a humidified incubator. 293T cells were used from passage 7 to 28, fibroblasts from passage 4 to 20 and urine-derived kidney cells from passage 14 to 20. When 70-90% confluent, cells were washed with Dulbecco’s Phosphate Buffered Saline (DPBS; Gibco, 14190-250) and passaged using 0.05% Trypsin (Gibco, 25300-054). Cells were cryo-preserved in liquid nitrogen in FBS with 10% DMSO (Sigma-Aldrich, D2438-50ML).

iPSCs were cultured in StemFlex media (Gibco, A3349401), in cell culture plates coated with an extracellular matrix hydrogel (R&D Systems, 3432-010-01) at 37°C and 5% CO_2_ in a humidified incubator^12^. iPSCs were thawed at passage 14 to 25 and used for not more than 10 passages. Cells were passaged with ReLeSR™ (STEMCELL Technologies, 100-0484) when 60-80 % confluent and kept in KnockOut Serum Replacement (Gibco, 10828028) with 10% DMSO and ROCK inhibitor (1:1000 from 10 mM stock, reconstituted in DMSO; STEMCELL Technologies, 72304) for long-term cryo-preservation in liquid nitrogen.

### 2.3. Generation of Knockout cells lines and prime editing

293T cells with knockouts of DLST, OGDH as well MMUT-KO1 and KO2 were previously generated^9^ and further characterized in this publication. CRISPR-Cas9 editing with homology directed repair was performed in 293T cells to generate MMUT-KO3 to 10 and MMUT-p.Y100C clones. Prime-editing was performed in 293T cells and iPSCs to introduce the variants MMUT-p.N219Y, MMUT-p.P615T and/or MMUT-p.R694W, using the PE3 system^22^. Experimental details are provided in Supporting Information.

### 2.4. Enzyme activity assay

MMUT enzyme activity assays were performed in crude cell lysis as previously described^23,24^ with recent modifications^25^. Data from primary fibroblasts was adapted from previously published data^9,21^.

### 2.5. Western Blotting

Western blots were performed as previously described^12^. Incubation with primary antibodies (MMUT: abcam, ab67869, 1:500; ACTB: Sigma-Aldrich, A1978, 1:1000) was performed overnight at 4°C while shaking and with the secondary antibody (anti-mouse horseradish peroxidase: SantaCruz, sc516102-cm, 1:5000) for 1 hour at room temperature. A detailed protocol can be found in Supporting Information.

### 2.6. Differentiation of iPSCs into neurons

iPSCs were differentiated into glutaminergic excitatory neurons using NGN2 - induced lentiviral programming, as previously described^26^. For details, see Supporting Information.

### 2.7. Extracellular flux analysis

The Mito Stress Test was performed on an XFp Analyzer or XFe96 Analyzer (Agilent) to measure oxygen consumption rates (OCR). Cells were seeded the day before the assay at a density of 12’500 cells/ well for 293T cells and 30’000 for fibroblasts, urine-derived epithelial kidney cells and iPSCs in 80 μl medium in the Seahorse XFe96/XF Pro cell culture microplates (Agilent, 103792-100). For 293T cells, plates were coated with poly-L-lysine (Sigma-Aldrich, P4707-50ML). A Countess 3 Automated Cell Counter (Thermo Fisher Scientific) with trypan blue stain (Invitrogen, T10282) was used for cell counting, following manufactures’ instructions. The differentiation of iPSCs was directly performed in the assay plates (see section 2.6). In detail, neuronal progenitors were seeded at day 3 of differentiation at a density of 100’000 cells/ well in PEI/rhLaminin-521 coated plates (0.05% PEI in borate buffer (Sigma-Aldrich, P3143); 1 μg/ml rhLaminin-521 (from 100 μg/ml stock in DPBS; Thermo Fisher Scientific, A29249)) and differentiated until day 14 or 21. Fresh medium (50 µl, containing 1 μg/ml rhLaminin-521) was added at least 1 h before assay preparation to ensure cell attachment.

On the day of the measurement, all cell types were washed twice with phenol-red free DMEM (Sigma-Aldrich, D5030), before media was exchanged to XF DMEM pH 7.4 (Agilent, 103575-100), supplemented with 2.0 g/l glucose (Gibco, A24940-01), 2 mM glutamine (Sigma-Aldrich, G7513) and 1 mM pyruvate (Gibco, 11360-070). For media stress conditions, concentrations of the supplements were adjusted accordingly, as described in the figure legends (galactose stock solution was prepared as 200 g/l solution in ddH_2_O; Sigma-Aldrich, G6404-25G). Cells were incubated at 37°C without CO_2_ for 60 min, before the measurement was started. Three or four cycles (3 min mixing and 3 min measuring) were recorded under basal conditions and after addition of oligomycin (Oligo; Sigma-Aldrich, O4876-25MG), carbonyl cyanide-p-trifluoromethoxyphenylhydrazone (FCCP; Sigma-Aldrich, C2920-10MG), and a combined injection of rotenone (Rot; Sigma-Aldrich, R8875-1G) and antimycin A (Anti A; Sigma-Aldrich, A8674-50MG). Concentrations were optimized for each cell type as followed: 293T cells: 2.0 μM Oligo, 1.5 μM FCCP and 0.5 μM Rot/Anti A; fibroblasts: 2.0 μM Oligo, 3.0 μM FCCP and 0.5 μM Rot/Anti A; urine-derived epithelial kidney cells: 2.0 μM Oligo, 1.0 μM FCCP and 0.5 μM Rot/Anti A; iPSCs, 1.5 μM Oligo, 0.5 FCCP and 0.5 μM Rot/Anti A; neurons: 2.0 μM Oligo, 1.5 μM FCCP, 1.0 μM Rot/Anti A. Inhibitors were dissolved in DMSO as 10 mM stock solutions and stored at -20°C or -80°C for FCCP. Integrity of the cell layer was manually checked after the assay, before media was completely removed and plates frozen at -80°C. Normalization by CyQUANT (Invitrogen, C7026) was used for 293T cells, fibroblasts, iPSCs and urine-derived epithelial kidney cells. Briefly, cells were lysed, and fluorescence was measured at 480/ 520 nm excitation/emission (Agilent, BioTek Synergy H1 Multimode Reader) after 5 min incubation at room temperature, protected from light.

Agilent Seahorse Wave Pro software (Agilent, Version 10.1.0.1) was used to analyze the data.

### 2.8. Mitochondrial fluorometric dyes

For 293T cells, 25’000 cells/well were seeded in black 96-well plates with clear bottom (Revvity, 6005225), coated with poly-L-lysine. The next day, cells were washed twice with phenol-red free DMEM (Thermo Fisher Scientific, A1443001) supplemented with 1 mM pyruvate, 2 mM glutamine, and 2.0 g/l glucose. Cell were incubated with 150 nM tetramethylrhodamine-methyl-ester perchlorate (TMRM; Sigma-Aldrich, T668) to assess mitochondrial membrane potential, 200 nM MitoTracker Green (Thermo Fisher Scientific, M7514) and 1 μg/ml Hoechst 33342 (Thermo Fisher Scientific, 62249) in phenol-red free DMEM for 30 min at 37°C. Rotenone (or DMSO as solvent control) was supplemented at 1 µM to induce dissipation of the mitochondrial membrane. Neurons at day 13 of differentiation were treated in the same way, with 100 nM TMRM and 100 nM MitoTracker Green. After incubation, dyes were diluted by replacing half of the medium with dye-free medium and directly measured on the Operetta CLS (Perkin Elmer). Nine images were taken per well, with 5% overlap. Harmony (Revvity, Version 4.9) software was used for automated analysis of each channel and calculation of TMRM/MitoTracker ratios. The analysis pipeline can be found in Supporting Information. Additional qualitative images of 293T cells were taken with an EVOS™ M3000 (Invitrogen).

### 2.9. Quantification of mtDNA/nDNA ratios

DNA from frozen cell pellets was isolated using the QIAmp DNA Mini kit (Qiagen, 51306). To quantify the relative mtDNA/nDNA ratios, a quantitative PCR was performed on a QUANTStudio7 (Thermo Fisher Scientific), using 10.2 ng/μl DNA and GoTaq^®^ qPCR Master Mix (2x; Promega, A600A). Relative ratios were calculated using the 2^−ΔΔCt^ formular. Primers against *ND1* were used for mitochondrial DNA^10^: fwd: 5’-ACACTAGCAGAGACCAACCG-3’, rev: 5’-GAAGAATAGGGCGAAGGGGC-3’; *ACTB* was used for nuclear DNA: fwd: 5’-TCACCCACACTGTGCCCATCTACGA-3’, rev: 5’-CAGCGGAACCGCTCATTGCCAATGG-3

### 2.10. Stable isotope tracing

Stable isotope tracing was performed as previously published for DLST-KO and OGDH-KO 293T cells^9^ and with optimizations for MMUT-KO and MMUT-p.N219Y 293T cells, as described in the Supporting Information.

### 2.11. Metabolite measurement by liquid chromatography-mass spectrometry

Metabolites were measured on an Orbitrap Astral Mass Spectrometer (Thermo Fisher Scientific), as previously described^27^ with modifications described in the Supporting Information.

### 2.12. Immunocytochemistry

Immunocytochemistry was performed on neurons grown on 8-well chamber slides (ibidi, 80841) and fixed with 4% paraformaldehyde (Electron Microscopy science, EMS-15710). Cells were permeabilized with 0.1% Triton X-100 (Sigma-Aldrich, X100-100ML, diluted in DPBS) for 7 min and quenched twice with 100 mM glycine for 10 min. Blocking was done with 1% BSA (Roche, 03117057001) in DPBS for 60 min at ambient temperature, before incubation with primary and secondary antibody (both diluted in DPBS). Nuclei were stained by incubation with Hoechst 33342 (Invitrogen, H3570; diluted 1:2’500 in DPBS) for 5 min at ambient temperature and slides were mounted with ProLong™ Diamond Antifade Mountant (Thermo Fisher Scientific, P36965) and covered with precision glass cover slips (No. 1.5H; Marienfeld, 7051-0107242). A list of antibodies, dilutions and incubation times can be found in Table S1. Images were acquired using an inverted Leica TCS SP8 confocal laser-scanning microscope equipped with 405-, 488-, 552-, and 638-nm solid-state diode lasers, with fluorophores excited using the appropriate laser lines and their emission recorded by two hybrid detectors over fluorophore-specific spectral ranges. High-resolution z-stacks were acquired with an HCX PL APO 63×/1.30 glycerol-immersion CS2 objective (Leica Microsystems), using a lateral pixel size of 85.9 nm, a z-step of 334.5 nm, 13–20 optical sections. Images were processed using the Imaris File Converter x64 (Bitplane AG/ Oxford Instruments, Version 11.0.0) and the Imaris x64 software (Oxford Instruments, Version 11.0.0).

### 2.13. Statistics and reproducibility

The performed statistical tests are described in the respective figure legends. GraphPad Prism (Version 10.1.2) was used for statistical analyses. All experiments were performed at least two to three times independently (if not indicated otherwise), and all datapoints are shown, with a clear description of what they represent (means of technical replicates were combined first, followed by combining experimental replicates). In this manuscript, we define biological replicates as individual clones with different genetic backgrounds, experimental replicates as repetitions of an experiment, and technical replicates as repeats of the same clone within one experiment. Schematics for Figures 4A, 5A and S1D were generated using biorender.com.

## 3. Results

### 3.1. MMUT-knockout cells retain normal mitochondrial energy production

Given previous findings of altered mitochondrial structure and function in MMUT-deficiency^9–13^, we investigated whether mitochondrial energy production via oxidative phosphorylation is consequently altered. Using CRISPR/Cas9 -based gene-editing in 293T cells, we generated ten individual MMUT-knockout (MMUT-KO) clones, each harboring either a premature stop-codon or frameshift causing deletion (Figure 1A). These occurred across the protein, including the N-terminal substrate binding domain and the C-terminal cofactor (AdoCbl) binding domain (Figure 1A). As controls (Ctrl), we utilized clones which had undergone the CRISPR/Cas9 process, but with no detectable edits. In contrast to Ctrl cells, all MMUT-KO clones had no detectable MMUT protein (Figure 1B) and undetectable MMUT activity (Figure 1C). In all cells, we examined the mitochondrial oxygen consumption rate (OCR) before and after application of OXPHOS inhibitors as an indicator of mitochondrial energy generation (see Methods). Overall, we found high clonal variability across both Ctrl and MMUT-KO clones (Figure S1A), but good reproducibility across experimental replicates (Figure S1B). Nevertheless, when all experiments and clones were combined, the OCR profiles of MMUT-KO and Ctrl cells showed a near-complete overlap, underscored by no difference in basal respiration, ATP production, maximal respiration and spare respiratory capacity (Figure 1D). Considering this finding, and to verify that the assay used is a robust method to detect disturbances in mitochondrial energy production, we examined 293T cells with a disturbed TCA cycle, namely knockouts of dihydrolipoamide succinyltransferase (DLST) or 2-oxoglutarate dehydrogenase (OGDH), both components of the 2-oxoglutarate dehydrogenase complex (OGDHc). Isolated DLST-KO and OGDH-KO clones showed reduced intracellular pool levels of malate and a trend for reduced (iso)citrate (Figure S1C). Furthermore, the normalized fractional atom contribution of ^13^C-glutamine to (iso)citrate was reduced and showed a trend to be reduced for malate (Figure S1D). Both findings are consistent with a functional disruption of the TCA cycle in OGDHc-KO cells. In these cells, we found a clearly reduced OCR profile and correspondingly lowered basal respiration, ATP production, maximal respiration and spare respiratory capacity (Figure S1E). Therefore, we are confident this assay and this cell-type are appropriate to assess mitochondrial energy production.

**Figure 1.**
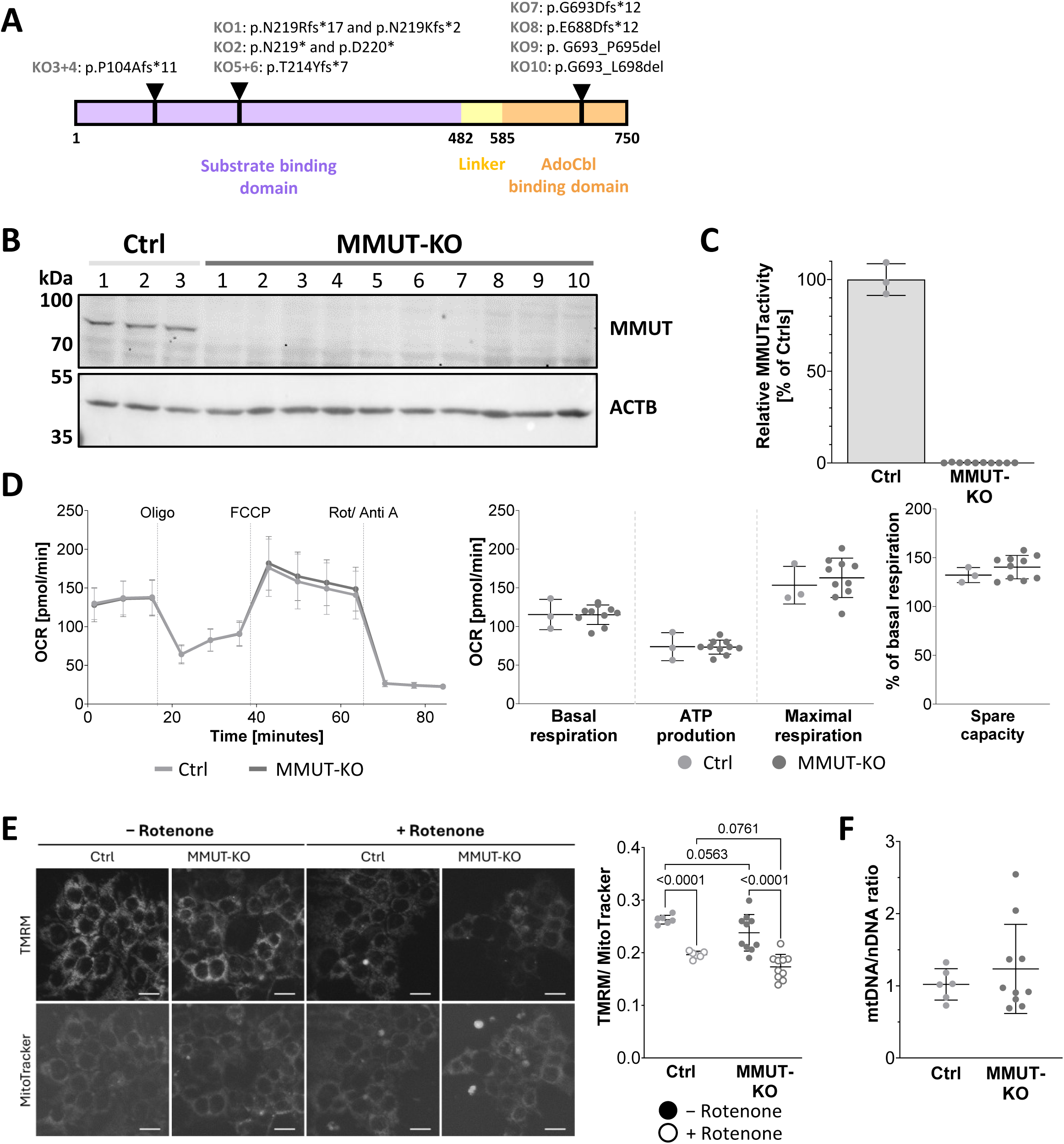
Mitochondrial respiration is not altered in 293T MMUT-KO. (A) Schematic overview of the location of genetic variants in MMUT-KO clones. (B) Western blot against MMUT in Ctrl (n=3) and MMUT-KO (n=10) 293T cells. ACTB was used as loading control. (C) Relative MMUT enzyme activity in Ctrl (n=3) and MMUT-KO (n=10) 293T cells. Data is normalized to the mean of unedited Ctrl cells (1649.7, 1441.7 and 1560.4 pmol/ mg/ min for experimental replicates). Each dot represents a biological replicate, based on the mean of three experimental replicates with two technical replicates. (D) Normalized OCR profiles and individual parameters of Ctrl (n=3) and MMUT-KO (n=10) cells. Data is shown as mean ± SD. Each dot represents a biological replicate, based on the mean of 2-4 technical replicates from eleven experimental replicates. Biological replicates were pooled for OCR profiles. (E) Left, representative images and right, quantification of TMRM and MitoTracker Green measurements in the presence and absence of rotenone (1 μM) in Ctrl (n=6) and MMUT-KO (n=10) cells. Data is shown as mean ± SD. Each dot represents a biological replicate, based on the mean of 4 technical replicates from two to twelve experimental replicates. A 2-way ANOVA with uncorrected Fisher’s LSD for multiple comparisons testing was used to compare genotypes and conditions. Scale bar is 20 µm. (F) Ratio of mitochondrial DNA (*ND1*) and nuclear DNA (*ACTB*) was determined by quantitative PCR on genomic DNA in Ctrl (n=6) and MMUT-KO (n=10). Data is shown as mean ± SD. Each dot represents a biological replicate, based on the mean of three technical replicates from one out of three representative experiments.

We consequently set out to determine if the lack of impact on mitochondrial bioenergetics in MMUT-KO cells reflected an overall conserved mitochondrial function. Using live cell imaging with the cell-permeable fluorescent dye tetramethylrhodamine methyl ester (TMRM), we found mitochondrial membrane potential to be similar between MMUT-KO and Ctrl cells, which was persistent when cells were treated with OXPHOS complex I inhibitor rotenone (Figure 1E). Likewise, quantitative PCR based estimation of mitochondrial abundance, found mitochondrial DNA (mtDNA) to nuclear DNA (nDNA) ratio to be unchanged between MMUT-KO and Ctrl cells (Figure 1F). Overall, these results suggest that the loss of MMUT does not result in altered mitochondrial energy production or mitochondrial homeostasis in 293T cells.

To verify if these findings are cell-type specific, we investigated primary skin-derived fibroblasts from individuals affected by MMUT-deficiency. We selected 8 cell lines harboring homozygous truncating variants (labelled MMUT-KO, Figure 2A) which we compared against five unaffected controls. As with 293T cells, MMUT-KO fibroblasts had undetectable protein levels (Figure 2B) as well as strongly reduced enzyme activity (Figure 2C). Nevertheless, as with 293T cells, we found no reduction in the OCR profiles, alongside unchanged basal respiration, ATP production, maximal respiration and spare respiratory capacity (Figure 2D). Together, these results suggest that complete loss of MMUT does not impact mitochondrial energy production in skin-derived fibroblasts either.

**Figure 2.**
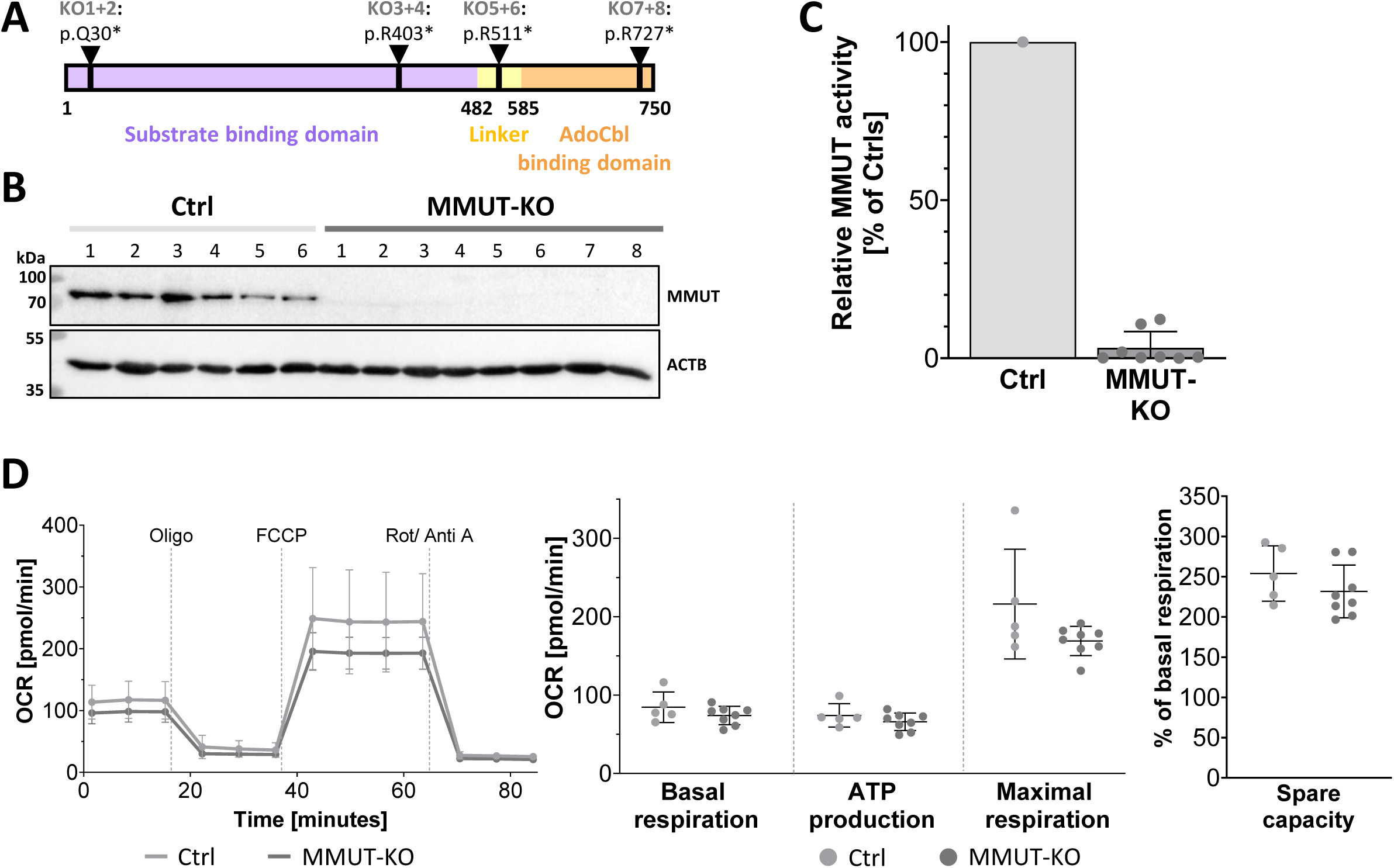
Mitochondrial respiration is not altered in fibroblasts with MMUT-KO from affected individuals. (A) Schematic overview of selected truncating variants. (B) Western blot against MMUT in Ctrl (n=6) and MMUT-KO (n=8) fibroblasts. ACTB was used as loading control. (C) Relative MMUT enzyme activity, compared to Ctrl (1’270 mol/ mg/ min). Data is shown as mean ± SD. Data points represent biological replicates, based on two technical replicates. (D) Normalized OCR profile and associated parameters in Ctrl (n=5) and MMUT-KO (n=8) fibroblasts. Data is shown as mean ± SD. Each dot represents a biological replicate, based on the mean of 3 - 4 technical replicates from two experimental replicates.

### 3.2. MMUT-p.N219Y causes decreased mitochondrial energy production in 293T cells

So far, 421 pathogenic or likely pathogenic variants in the *MMUT* gene, including 168 missense variants, have been described^28^ (Figure S2A). To investigate whether nonsense variants, which lead to a complete lack of protein, and missense variants, which can have variable effects on protein levels, differ in their impact on mitochondrial energy production, we selected four variants for deeper characterization: MMUT-p.Y100C, MMUT-p.N219Y, MMUT-p.P615T and MMUT-p.R694W (Figure 3A). These variants have been frequently detected in a homozygous state in a European cohort^21^, cover both mut^−^ (residual activity and cobalamin responsive^29^) and mut^0^ (undetectable activity^29^) subtypes^30–32^, and are inserted in the substrate binding domain and cofactor binding domain of MMUT^21^. In 293T cells, we generated three homozygous clones of each variant using prime editing (see Methods and Supporting Information). Cells harboring these variants had unchanged (MMUT-p.Y100C), decreased (MMUT-p.N219Y, MMUT-p.R694W) or undetectable protein levels (MMUT-p.P615T) (Figure 3B) along with strongly reduced to undetectable residual enzyme activity (Figure 3C). Examination of mitochondrial OCR revealed that only MMUT-p.N219Y clones had an altered OCR profile, with reduced spare respiratory capacity (Figure 3D). In addition, using an ordinary one-way ANOVA, we found that maximal respiration was not the same for all MMUT variants (p = 0.0427), but multiple testing for specific comparisons to Ctrl cells did not reach significance (p = 0.0762 for MMUT-p.N219Y). However, individual p-values showed a significant decrease in maximal respiration and a trend in decreased ATP production for MMUT-p.N219Y clones (Figure 3D), making this variant a promising candidate for decreased mitochondrial energy production and further investigations. These results are in line with the altered mitochondrial membrane potential in clones harboring the MMUT-p.N219Y variant (Figure 3E and Figure S2B). However, mtDNA/nDNA ratios were unchanged in these cells (Figure 3F).

**Figure 3:**
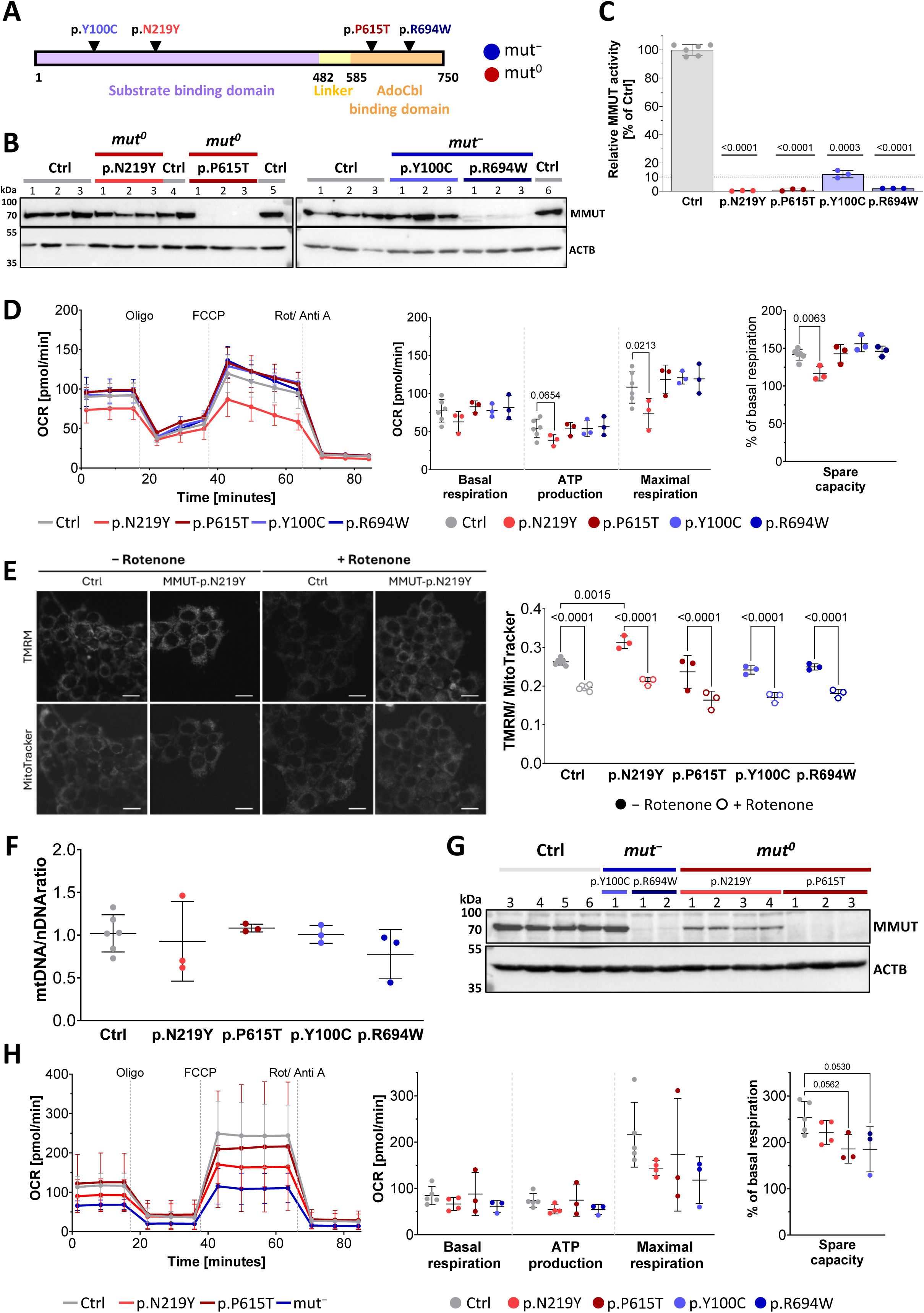
MMUT missense variants show decreased mitochondrial energy production in 293T cells and fibroblasts. (A) Schematic overview of selected MMUT-missense variants. Blue tones represent *mut^−^*, red tones represent *mut*^0^, and grey for control. (B) Western blot against MMUT in Ctrl (n=6) and four MMUT missense variants (n=3 each) in 293T cells. ACTB was used as loading control. (C) Relative MMUT enzyme activity in Ctrl (n=6) and four MMUT missense variants (n=3 each) in 293T cells. Data is normalized to the mean of all Ctrl cells (1165.4, 1089.7 and 1123.0 pmol/ mg/ min for experimental replicates). Each dot represents a biological replicate, based on the mean of three experimental replicates with two technical replicates. 10% residual activity are indicated by a dotted line Values across experiments were normalized as a fraction of the average of controls in each experiment and tested to be different to 100% based on a one-sample t-test. (D) Normalized OCR profiles and individual parameters of Ctrl (n=6) and MMUT missense variants (n=3 each) in 293T cells. Data is shown as mean ± SD. Each dot represents a biological replicate, based on the mean of 2-6 technical replicates from two experimental replicates. Biological replicates were pooled for OCR profiles. An ordinary one-way ANOVA with Dunnett’s multiple comparison was used to compare each variant to Ctrl cells. The one-way ANOVA was significant for maximal respiration, but Dunnett’s multiple comparison test was not significant (p=0.0762). Individual p-values, calculated by ordinary one-way ANOVA with two-stage linear step-up procedure of Benjamini, Krieger and Yekutieli, are shown in the graph. Adjusted p-value (one-way ANOVA following Dunnett’s multiple comparison test) is reported for spare respiratory capacity. (E) Left, representative images for one Ctrl and one MMUT-p.N219Y clone and right, quantification of TMRM and MitoTracker Green measurements in the absence (closed circle) and presence (open circle) of rotenone (1 μM) in Ctrl (n=6) and MMUT missense variants (n=3 each) in 293T cells. Scale bar is 20 µm. Data is shown as mean ± SD. Each dot represents a biological replicate, based on the mean of 4 technical replicates from two to twelve experimental replicates. A two-way ANOVA with Tukey’s multiple comparisons testing was used to compare genotypes and conditions. (F) Ratio of mitochondrial DNA (ND1) and nuclear DNA (ACTB) was determined by quantitative PCR on genomic DNA in Ctrl (n=6) and four MMUT missense variants (n=3 each). Data is shown as mean ± SD. Each dot represents a biological replicate, based on the mean of three technical replicates from one out of three representative experiments. (G) Western blot against MMUT in Ctrl (n=4) and affected fibroblast lines (n=10). ACTB was used as loading control. (H) Normalized OCR profiles and individual parameters in Ctrl (n=5) and MMUT-deficient (n=10 in total) fibroblasts. Data is shown as mean ± SD. Each dot represents a biological replicate, based on the mean of 3-4 technical replicates from two experimental replicates. Biological replicates were pooled for OCR profiles. Adjusted p-value (one-way ANOVA following Dunnett’s multiple comparison test) is reported for spare respiratory capacity.

To determine if this finding was conserved across cell-types, we identified skin-derived fibroblasts from individuals affected by MMUT-deficiency, expressing the same variants in a homozygous state^9,24^. As with 293T cells, MMUT-p.Y100C had unchanged, MMUT-p.N219Y and p.R694W decreased, and MMUT-p.P615T had undetectable protein levels (Figure 3G), while all cell lines had strongly reduced or undetectable residual enzyme activity (Figure S2C). Investigation of mitochondrial OCR suggested that all cells harboring MMUT missense variants had an altered OCR profile (Figure 3H). However, although numerically decreased, MMUT-p.N219Y was not significantly different from Ctrl cells in any individual parameter, while interestingly, cell lines harboring the mut^−^ variants MMUT-p.Y100C and p.R694W had a trend for reduced spare respiratory capacity (Figure 3H). These results confirm that the p.N219Y variant of MMUT impacts mitochondrial energy production in 293T cells and suggests the presence of variant specific and cell-type specific effects on mitochondrial homeostasis and energy production.

### 3.3. Altering TCA cycle fuel sources exacerbates the disturbance of mitochondrial energy generation in MMUT-p.N219Y expressing cells

So far, the presented results were conducted under media conditions containing supraphysiological concentrations of glucose, glutamine and pyruvate, which all fuel the TCA cycle (Figure 4A). However, and in line with previously published results^9^, alongside increased levels of methylmalonic acid (Figure S3A), we found pool levels of glutamine to be increased in 293T MMUT-p.N219Y clones compared to controls (Figure S3B), but normalized fractional atom contribution of ^13^C-glutamine into TCA cycle metabolites was unchanged (Figure S3C). This suggests that MMUT-deficient cells may use alternative TCA cycle sources (e.g. glucose or pyruvate) to fuel mitochondrial OXPHOS, thus masking a potential defect in mitochondrial energy production. Therefore, we examined the impact of depletion or sole provision of each of the three fuel sources for both MMUT-KO and MMUT-p.N219Y clones compared to Ctrl cells and standard conditions (Figure 4B and Figure S4A). In addition, galactose was provided as an alternative TCA cycle fuel source, which has previously been shown to exacerbate mitochondrial energetic deficits by limiting glycolysis and forcing cells to utilize OXPHOS, fueled by glutamine and pyruvate (Figure 4A)^33^.

**Figure 4.**
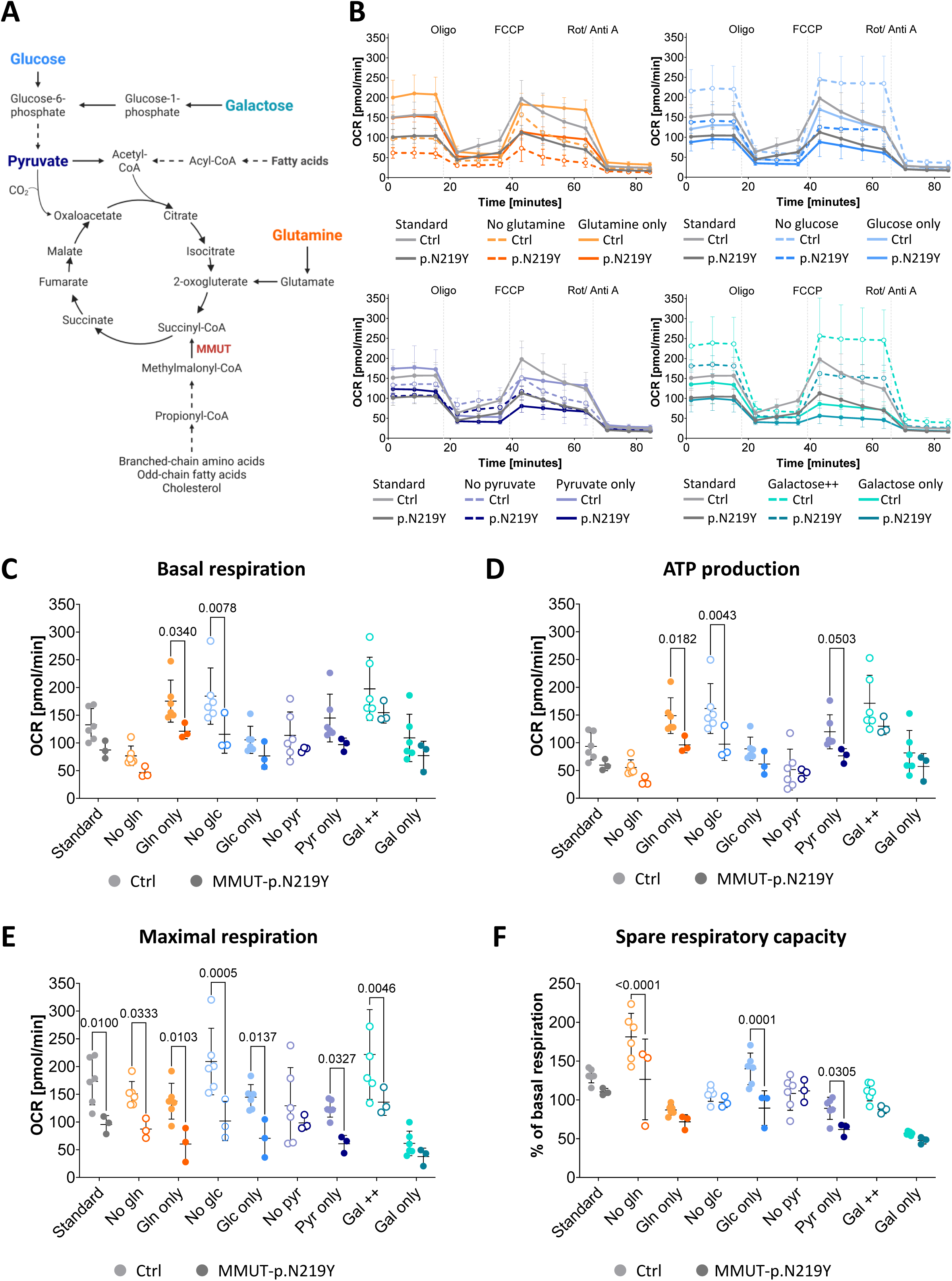
Examination of TCA cycle fuel sources on mitochondrial energy production in MMUT-p.N129Y 293T cells. (A) Schematic overview of TCA cycle fuel sources. (B) Normalized OCR profiles in Ctrl (n=6) and MMUT-p.N219Y (n=3) 293T cells under standard conditions (2.0 mM glutamine, 2.0 g/l glucose, 1.0 mM pyruvate) and stress conditions (0.25 mM glutamine in glutamine only; 0.25 g/l glucose for glucose only; 0.125 mM pyruvate for pyruvate only; 2.0 g/l galactose for galactose++ and 0.25 g/l for galactose only. Colors refer to TCA cycle fuel sources depicted in A, with light shades for Ctrl and darker shades for MMUT-p.N219Y clones. Biological replicates were pooled for OCR profiles, and profiles for standard conditions are pooled from four experiments. (C-F) Normalized parameters under standard and stress conditions. Each dot represents one biological replicate based on the mean of 2-4 technical replicates, or the mean of four experimental replicates with 2-4 technical replicates for standard conditions. A two-way ANOVA with Tukey’s multiple comparisons test was used to compare the variant effect across all media conditions, reporting the significant adjusted p-values (p < 0.05). Abbreviations: Gln: glutamine; Glc: glucose; Pyr: pyruvate; Gal: galactose.

Basal respiration was reduced in both MMUT-KO and Ctrl cells when glutamine was depleted or glucose was the sole fuel source, whereas it increased when glucose was replaced with galactose and trended to increase when glutamine was the sole supplemented source (Figure S4A and S4B). Notably, in all conditions, no differences between MMUT-KO and Ctrl cells were detected (Figure S4). Importantly, although cells expressing MMUT-p.N219Y showed the same overall responses to altered media composition (Figure 4B), compared to Ctrl cells, basal respiration was reduced with glutamine as the sole source and in media without glucose (Figure 4C).

Changes in ATP production were in line with changes in basal respiration in MMUT-KO and Ctrl cells: compared to standard media conditions, glutamine depletion decreased ATP production, while sole glutamine, pyruvate and galactose supplementation and glucose depletion increased ATP production (Figure S4C). Cells expressing MMUT-p.N219Y showed these same responses. However, compared to Ctrl cells they additionally showed reduced ATP production when glutamine was the only source added or glucose was depleted (Figure 4D). Furthermore a trend for decreased ATP production was found when pyruvate was the sole source (Figure 4D).

For both MMUT-KO and Ctrl cells, compared to standard conditions, maximal respiration was decreased in all conditions that lacked pyruvate (i.e. glutamine only, glucose only, no pyruvate, and galactose only) (Figure S4D). In addition to this overall response, compared to control cells, MMUT-p.N219Y cells had reduced maximal respiration in almost all tested conditions (excepting: no pyruvate and galactose only), including standard conditions (Figure 4E).

Finally, spare respiratory capacity was similarly reduced in almost all media stress conditions compared to standard conditions in MMUT-KO and Ctrl cells, except it was increased upon glutamine depletion and unchanged when glucose was the sole source (Figure S4E). Again, compared to Ctrl cells, those harboring the MMUT-p.N219Y variant showed decreased spare respiratory capacity under glutamine depletion and sole supplementation with glucose or pyruvate (Figure 4F).

Overall, the changes in media composition did not induce a phenotype in MMUT-KO clones, but indeed exacerbated the phenotype of MMUT-p.N219Y 293T cells.

### 3.4. Decreased mitochondrial energy production in MMUT-deficient epithelial kidney cells, but not induced pluripotent stem cells nor neurons

The brain and kidneys are two of the most severely affected organs in MMA, potentially due to their high energy demand^2,5,12^. Thus, we investigated how mitochondrial energy production is altered in cells representing these tissues.

We first examined urine-derived epithelial kidney cells^20^ harboring heterozygous MMUT missense variants p.S288P and p.H386R (both classified as *mut*^0^ ^30,34^). We identified decreased protein levels in all affected cell lines (Figure S5A). Consistent with previous publications^10,17^, examination of mitochondrial OCR revealed decreased basal respiration, ATP production, and maximal respiration in these cells (Figure S5B).

Next, we examined our previously described induced pluripotent stem cells (iPSC) derived from fibroblasts of two affected individuals and two unaffected controls ^12^. Additionally, we used prime editing to introduce the MMUT-p.N219Y variant in one unaffected iPSC line, resulting in two further MMUT-deficient clones and one unedited isogenic control (Figure 5A). As expected, all iPSC lines harboring the MMUT-p.N219Y variant showed reduced MMUT protein levels compared to Ctrl cells (Figure 5B), and nearly undetectable enzyme activity (Figure 5C). However, basal respiration, ATP production, and maximal respiration were unchanged compared to Ctrl cells under standard conditions, while remarkably, spare respiratory capacity was increased in MMUT-p.N219Y clones compared to control cells (Figure 5D). To determine if this was an iPSC specific effect, we assayed the two fibroblast lines from which the affected iPSC lines were derived, and found they had no change in OCR parameters, including spare respiratory capacity (Figure S6).

**Figure 5.**
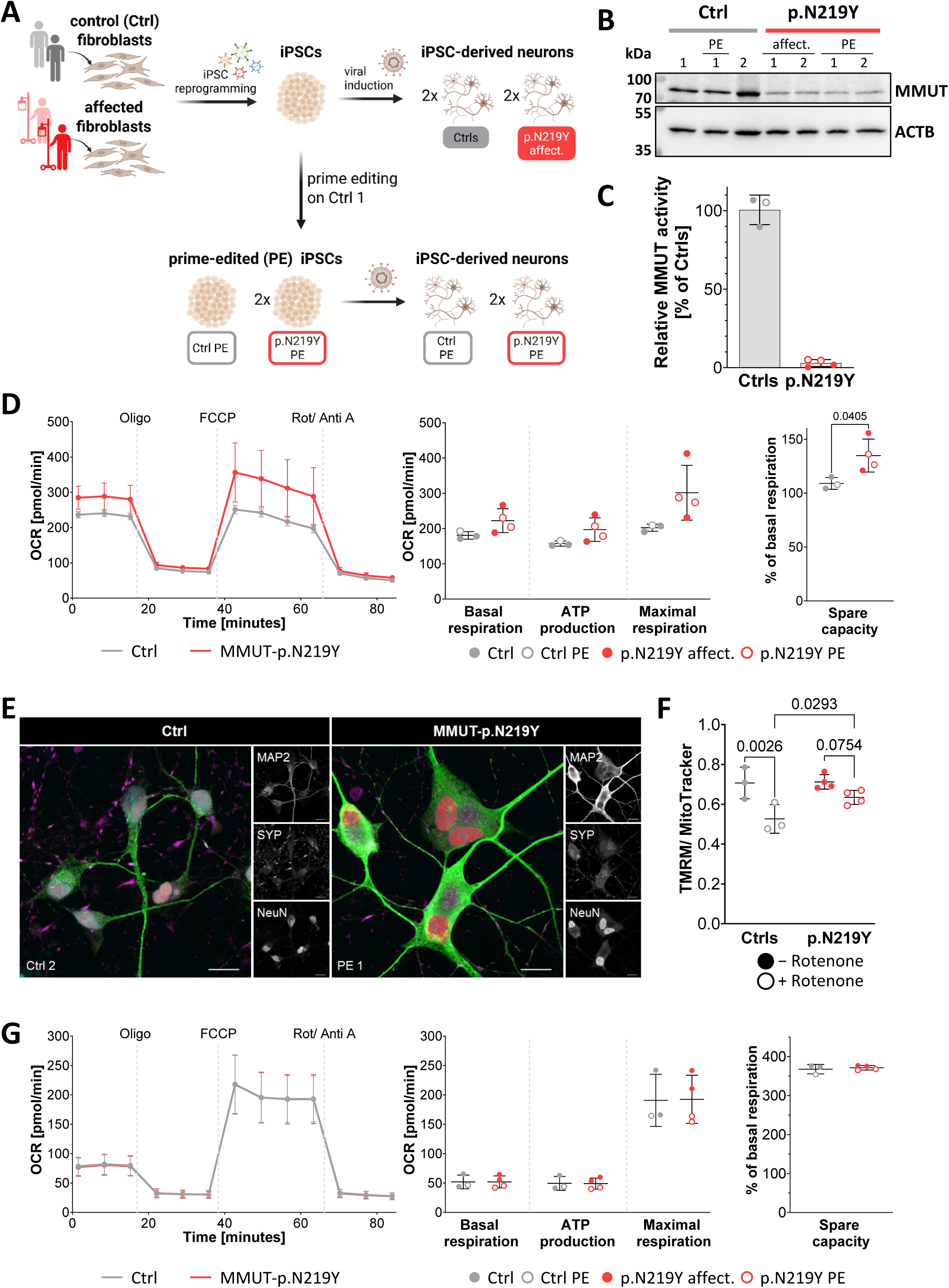
Mitochondrial energy production in induced pluripotent stem cells and neurons. (A) Schematic overview of the origin and generation of the iPSC and neuronal cell lines. Abbreviations: affect. = Affected cell line; PE = prime edited. (B) Western blot against MMUT in Ctrl (n=3) and MMUT-p.N219Y (n=4) clones in iPSCs. ACTB was used as loading control. Uncropped images of all membranes can be found in Figure S7. (C) Relative MMUT enzyme activity in Ctrl (n=3) and MMUT-p.N219Y clones (n=4) in iPSCs. Data is normalized to the mean of all Ctrl cells (416.3 pmol/ mg/ min). Each dot represents a biological replicate, based on the mean of three experimental replicates with two technical replicates. (D) Normalized OCR profiles and individual parameters of Ctrl (n=3) and MMUT-p.N219Y clones (n=4) in iPSCs. Data is shown as mean ± SD. Each dot represents a biological replicate, based on the mean of 10-12 technical replicates from three experimental replicates. Biological replicates were pooled for OCR profiles. An unpaired t-test was applied. (E) Immunocytochemistry for neurons at day 14 of differentiation in one Ctrl and one MMUT-p.N219Y PE line. Shown are microtuble associated protein 2 (MAP2, green), Synaptophysin (SYP, magenta), NeuN (red) and nuclei stained with Hoechst (cyan). Scale bar is 10 µm. Images of the other cell lines are shown in Figure S7A. (F) Quantification of TMRM and MitoTracker Green measurements in the presence and absence of rotenone (1 μM) in Ctrl (n=3) and MMUT-p.N219Y (n=4) in neurons at day 13 of differentiation. Data is shown as mean ± SD. Each dot represents a biological replicate, based on the mean of 2-6 technical replicates. A 2-way ANOVA with uncorrected Fisher’s LSD for multiple comparisons testing was used to compare genotypes and conditions. (G) OCR profiles and individual parameters of Ctrl (n=3) and MMUT-p.N219Y clones (n=4) in neurons at day 14 of differentiation. Data is shown as mean ± SD. Each dot represents a biological replicate, based on the mean of 2-11 technical replicates from three experimental replicates. Biological replicates were pooled for OCR profiles.

To examine the impact of this variant in neuronal cells, lentiviral-induced forward programming was used for the differentiation of iPSCs into excitatory neurons^26^. Neuronal cell fate was confirmed by the presence of microtubule-associated protein 2 (MAP2), synaptophysin (SYP) and NeuN, all markers for mature neurons^35^, expressed on day 14 of differentiation (Figure 5E and S7A). We found no difference in the mitochondrial potential between Ctrl and MMUT-p.N219Y neurons in the absence of rotenone, however when exposed to rotenone MMUT-p.N219Y neurons had a comparatively increased TMRM/MitoTracker ratio due to a lack of response to this complex I inhibitor (Figure 5F). Examination of mitochondrial OCR in neurons (on day 14 of differentiation) revealed no difference between MMUT-p.N219Y and Ctrl cell lines (Figure 5G). Prolonging the differentiation until day 21 resulted in the same outcome (Figure S7B). Together, these results confirm that urine-derived epithelial kidney cells have reduced mitochondrial energy production, but this defect is not found in iPSCs nor their *in vitro* derived neurons.

## 4. Discussion

In this study, we systematically investigated the consequences of MMUT-deficiency on mitochondrial energy production using extracellular flux analysis, a method widely used to assess mitochondrial function^19^. We found variant-dependent changes within a given cell type, as well as cell-type-dependent changes specific to the MMUT-p.N219Y variant.

Whereas basal respiration was unaffected in all tested cell types apart from urine-derived epithelial kidney cells, maximal respiration and spare respiratory capacity were more frequently reduced. The ability to maintain a bioenergetic profile with spare respiratory capacity is considered to be essential for a healthy mitochondrial network^30,31^. Thus, our findings suggest that MMUT-deficient cells are more susceptible to stress^32^, induced by the disruption of the proton gradient by the uncoupling protonophore FCCP. Importantly, the observed differences between cell types must be considered against the background of substantial metabolic differences and mitochondrial heterogeneity across tissues and cell-types, which have been widely reported^39,40^.

Furthermore, we showed that stress conditions induced by depletion or sole supplementation of TCA cycle sources exacerbated the decrease in mitochondrial energy production in MMUT-p.N219Y 293T cells. In particular, conditions that forced a shift from glycolysis to OXPHOS (i.e. no glucose in the medium) caused an decrease in basal respiration and ATP production. Galactose as alterative carbohydrate source induces this shift as well, which is mainly driven by the usage of alternative TCA cycle fuel sources, including glutamine, pyruvate, fatty acids and amino acids^33^. Here, we investigated the acute response to changes in media conditions, thus the sole supplementation with galactose had no effect on basal respiration, as the conversion of galactose into glucose-1-phosphate is rather slow^41^ and requires transcriptional changes for an efficient processing^42,43^. In addition, depletion of pyruvate caused a decrease in spare respiratory capacity, as previously reported^31,33,34^. However, apart from these specific findings, the same overall responses to altered media conditions were observed for MMUT-p.N219Y, MMUT-KO and Ctrl 293T cells, suggesting that all cells were able to rapidly adapt to changes in the media composition.

The accumulation of potentially toxic intermediates^46^ and secondary metabolites, such as ammonium and lactate^47,48^, is reported to cause mitochondrial dysfunction in MMA by inhibiting TCA cycle enzymes and OXPHOS complexes^49–51^, ultimately causing an increase in reactive oxygen species, which further induces mitochondrial damage^4^. We found elevated levels of methylmalonic acid in MMUT-p.N219Y 293T cells. Nonetheless, this cannot fully explain the decrease in OCR in these cells, because MMUT-KO 293T cells had elevated levels of methylmalonic acid as well, but had normal OCR. In line with this observation, previous studies showed varying effects of methylmalonic acid on respiratory chain complexes when administered *in vitro* to rat tissues^51–55^ and human neuroblastoma SH-SY5Y cells^56^. Additionally, methylmalonic acid had no inhibitory effect on mitochondrial respiration in murine muscle tissue^57^.

While nonsense variants will most likely result in the full absence of the enzyme, the consequences of missense variants are harder to predict as multiple factors may be affected, including protein expression, protein stability, cellular localization, residual activity and degradation^58^. The described differences in mitochondrial homeostasis and energy production between cells harboring MMUT-KO and MMUT missense variants suggest that additional cellular mechanisms might be responsible for inducing or compensating for mitochondrial dysfunction in MMUT-deficient cells. One possible compensatory mechanism is transcriptional adaptation, which describes the upregulation of alternative genes mediated by nonsense mediated decay (NMD). NMD is specifically triggered by premature stop codons in mRNA transcripts carrying nonsense variants^59–63^, as shown in a study about Duchenne muscular dystrophy^64^. Similarly, it has been reported for Marfan syndrome^65^ and inherited cardiomyopathies^66^ that missense variants can have more detrimental effects than nonsense variants. However, such an observation has not been reported for MMA. Paradoxically, mut^−^ variants, which are more often missense variants^21,67,68^, correlate with milder symptoms and later disease onset^69–71^. Thus, more studies are necessary to better understand genotype-phenotype correlations in MMA and how *in vitro* findings can be translated *in vivo*.

Apart from altered mRNA degradation, nonsense and missense variants may also differentially disrupt protein-protein interactions. MMUT is known to form a complex with methylmalonic aciduria type A protein (MMAA, MIM: #607481)^72,73^ and to interact with multiple enzymes from the TCA cycle^9^. While most protein-protein interactions may be lost for nonsense variants, missenses variants may show more complex interaction defects, including loss of all interactions, allosteric effects and edgetic effects, which cause the loss of a specific subset of interactions^74–76^. Altered protein-protein interactions are hypothesized to be linked to different disease phenotypes and severity^75–77^ as for example reported for osteogenesis imperfecta^75^. Finally, it has been reported that MMA-causing variants in the adenosylcobalamin processing enzyme MMAA, cause the loss of interaction with MMUT^78^, thus it is likely that this is also true, when MMUT is harboring a missense variant.

Mitochondrial dysfunction is commonly associated with impaired mitochondrial energy production^79–82^. Our results show that MMUT-deficiency does not necessarily affect mitochondrial energy production, despite evidence for disruptions in mitochondrial homeostasis and previous reports of mitochondrial dysfunction^10–12,17^.

The central question remaining is how the cells in our study maintain mitochondrial homeostasis. One possible explanation is that MMUT dysfunction is compensated by other processes, most likely in a cell-type- and variant-dependent manner.

By using different cell types, including primary cells derived from affected individuals, as well as a neuronal cell model for otherwise hardly accessible tissues, and different types of MMUT variants in combination with their respective isogenic controls, we provide a broad and systematic overview of how mitochondrial energy production is affected in MMA. However, our study has important limitations. We predominately used one method to investigate mitochondrial dysfunction and all experiments were performed *in vitro*. Furthermore, the iPSC-derived neurons were measured rather early in their differentiation^83,84^ and we focused only on the missense variant MMUT-p.N219Y, for which we previously reported mitochondrial dysfunction^12^. Additional experiments, e.g. via transcriptomic and proteomic analyses, may be necessary to explore regulatory mechanisms, both across cell-types and variants. Clinically, both truncating variants (i.e. MMUT-KO) and the missense variant MMUT-p.N219Y are classified as *mut*^0^ and frequently result in severe disease with overlapping symptoms^21,70,85,86^, so it is not clear how the different regulatory mechanisms found here may impact phenotypic findings. With that in mind, determining how rescue of energy production is achieved *in vitro* and can be translated *in vivo*, could build the foundation for potential new therapeutic strategies for MMA.

## 5. Author contribution

M.A.G., D.S.F. and M.R.B. contributed to the conception and design of the study.

M.A.G. carried out the experimental work contributing to the majority of the paper.

M.A.G. and D.S.F. prepared the manuscript with input from all other authors. P.C.M. cultured and differentiated the iPSCs into neurons, performed ICC and wrote parts of the method section. L.T. performed LCMS experiments and wrote parts of the method section. C.T.G. contributed to generating MMUT-KO cell lines. J.L. contributed to OCR measurements of fibroblasts and Western blots of urine-derived epithelial kidney cells. M.C.S.D. generated the iPSC lines. C.B. generated MMUT-KO and MMUT-variants in 293T cells and iPSCs. S.L. performed MMUT activity assays. A.S., S.K. and O.D. provided the urine-derived epithelial kidney cells. R.J.M. contributed to analysis and interpretation of LCMS experiments. F.T. contributed to cell maintenance and performed ^13^C-glutamine labelling experiments. All authors contributed to the manuscript and approved the submitted version.

## 6. Acknowledgements

The authors thank Vito R.T. Zanotelli for his advice with statistical analyses, James A. Crowe for his support with the optimization and analysis of high-throughput imaging experiments with TMRM and MitoTracker in 293T cells and neurons and Stephan Wüest for providing access to the XFPro Analyzer. Imaging was performed with equipment maintained by the Center for Microscopy and Image Analysis, University of Zurich. M.A.G., C.T.G., L.T., O.D., D.S.F. and M.R.B. are supported by the University Research Priority Program of the University of Zurich (URPP) ITINERARE – Innovative Therapies in Rare Diseases. We acknowledge support by the Swiss National Science Foundation to R.J.M [221617 and 10007828], D.S.F. [219127 and 239897] and M.R.B. [212505]. P.C.M is supported by the Anna Mueller Grocholski-Stiftung. This work was supported by the UMZH platform Zurich Trace.

## 7. Conflict of interest

The authors declare that they have no conflict of interest.

## 8. Data availability statement

The data that supports the findings of this study are available on request.

## Notes

### Competing Interest Statement

The authors have declared no competing interest.

## 9#References

1. Froese, D. S. & Gravel, R. A. Genetic disorders of vitamin B_12_ metabolism: eight complementation groups – eight genes. Expert Rev. Mol. Med. 12, e37 (2010).

2. Forny, P. et al. Guidelines for the diagnosis and management of methylmalonic acidaemia and propionic acidaemia: First revision. J. Inherit. Metab. Dis. 44, 566–592 (2021).

3. Ballhausen, D., Mittaz, L., Boulat, O., Bonafé, L. & Braissant, O. Evidence for catabolic pathway of propionate metabolism in CNS: expression pattern of methylmalonyl-CoA mutase and propionyl-CoA carboxylase alpha-subunit in developing and adult rat brain. Neuroscience 164, 578–587 (2009).

4. Du, M. et al. Metabolic toxicity and neurological dysfunction in methylmalonic acidemia: from mechanisms to therapeutics. Mol. Med. 31, 333 (2025).

5. Haijes, H. A., Jans, J. J. M., Tas, S. Y., Verhoeven-Duif, N. M. & Van Hasselt, P. M. Pathophysiology of propionic and methylmalonic acidemias. Part 1: Complications. J. Inherit. Metab. Dis. 42, 730–744 (2019).

6. Manoli, I. et al. Targeting proximal tubule mitochondrial dysfunction attenuates the renal disease of methylmalonic acidemia. Proc. Natl. Acad. Sci. 110, 13552–13557 (2013).

7. Ramon, C., Traversi, F., Bürer, C., Froese, D. S. & Stelling, J. Cellular and computational models reveal environmental and metabolic interactions in *MMUT* -type methylmalonic aciduria. J. Inherit. Metab. Dis. 46, 421–435 (2023).

8. Anzmann, A. F. et al. Multi-omics studies in cellular models of methylmalonic acidemia and propionic acidemia reveal dysregulation of serine metabolism. Biochim. Biophys. Acta BBA - Mol. Basis Dis. 1865, 165538 (2019).

9. Forny, P. et al. Integrated multi-omics reveals anaplerotic rewiring in methylmalonyl-CoA mutase deficiency. Nat. Metab. 5, 80–95 (2023).

10. Luciani, A. et al. Impaired mitophagy links mitochondrial disease to epithelial stress in methylmalonyl - CoA mutase deficiency. Nat. Commun. 11, 970 (2020).

11. Chandler, R. J. et al. Mitochondrial dysfunction in *mut* methylmalonic acidemia. FASEB J. 23, 1252–1261 (2009).

12. Denley, M. C. S. et al. Mitochondrial dysfunction drives a neuronal exhaustion phenotype in methylmalonic aciduria. *Commun*. Biol. 8, 410 (2025).

13. Schumann, A. et al. The impact of metabolic stressors on mitochondrial homeostasis in a renal epithelial cell model of methylmalonic aciduria. Sci. Rep. 13, 7677 (2023).

14. Zsengellér, Z. K. et al. Methylmalonic acidemia: A megamitochondrial disorder affecting the kidney. Pediatr. Nephrol. 29, 2139–2146 (2014).

15. Head, P. E. et al. Aberrant methylmalonylation underlies methylmalonic acidemia and is attenuated by an engineered sirtuin. Sci. Transl. Med. 14, eabn4772 (2022).

16. De Keyzer, Y. et al. Multiple OXPHOS Deficiency in the Liver, Kidney, Heart, and Skeletal Muscle of Patients With Methylmalonic Aciduria and Propionic Aciduria. Pediatr. Res. 66, 91–95 (2009).

17. Köpfer, F. et al. Effects of anserine on oxidative stress and on cell barrier integrity in methylmalonic aciduria. Sci. Rep. 15, 32933 (2025).

18. Stanescu, S. et al. Mitochondrial dysfunction in methylmalonic acidemia: A pilot study using Seahorse technology in peripheral blood. Mol. Genet. Metab. Rep. 45, 101251 (2025).

19. Yoo, I., Ahn, I., Lee, J. & Lee, N. Extracellular flux assay (Seahorse assay): Diverse applications in metabolic research across biological disciplines. Mol. Cells 47, 100095 (2024).

20. Ruppert, T. et al. Molecular and biochemical alterations in tubular epithelial cells of patients with isolated methylmalonic aciduria. Hum. Mol. Genet. ddv405 (2015) doi:10.1093/hmg/ddv405.

21. Forny, P. et al. Molecular Genetic Characterization of 151 *Mut* -Type Methylmalonic Aciduria Patients and Identification of 41 Novel Mutations in *MUT*. Hum. Mutat. 37, 745–754 (2016).

22. Anzalone, A. V. et al. Search-and-replace genome editing without double-strand breaks or donor DNA. Nature 576, 149–157 (2019).

23. Baumgartner, R. in The Cobalamins, Methods in Hematology (Churchill Livingstone, 1983).

24. Causey, A. G. & Bartlett, K. A radio-HPLC assay for the measurement of methylmalonyl-CoA mutase. Clin. Chim. Acta 139, 179–186 (1984).

25. Forny, P., Froese, D. S., Suormala, T., Yue, W. W. & Baumgartner, M. R. Functional Characterization and Categorization of Missense Mutations that Cause Methylmalonyl- C o A Mutase ( MUT ) Deficiency. Hum. Mutat. 35, 1449–1458 (2014).

26. Zhang, Y. et al. Rapid Single-Step Induction of Functional Neurons from Human Pluripotent Stem Cells. Neuron 78, 785–798 (2013).

27. Cherkaoui, S. et al. Reprogramming neuroblastoma by diet-enhanced polyamine depletion. Nature 646, 707–715 (2025).

28. National Center for Biotechnology Information (NCBI). ClinVar.Bethesda (MD): National Library of Medicine (US). ClinVar Search Results: MMUT [gene] AND (pathogenic [clinical-significance] OR likely pathogenic [clinical-significance]). (2026).

29. Willard, H. F. & Rosenberg, L. E. Inherited Methylmalonyl CoA Mutase Apoenzyme Deficiency in Human Fibroblasts. J. Clin. Invest. 65, 690–698 (1980).

30. Lempp, T. J. et al. Mutation and biochemical analysis of 19 probands with mut0 and 13 with mut− methylmalonic aciduria: Identification of seven novel mutations. Mol. Genet. Metab. 90, 284–290 (2007).

31. Acquaviva, C. et al. N219Y, a new frequent mutation among mut° forms of methylmalonic acidemia in Caucasian patients. Eur. J. Hum. Genet. 9, 577–582 (2001).

32. Peters, H. L. et al. Molecular studies in mutase-deficient (MUT) methylmalonic aciduria: identification of five novel mutations. Hum. Mutat. 20, 406–406 (2002).

33. Pinho, S. A. et al. Mitochondrial and metabolic remodelling in human skin fibroblasts in response to glucose availability. FEBS J. 289, 5198–5217 (2022).

34. Worgan, L. C. et al. Spectrum of mutations in *mut* methylmalonic acidemia and identification of a common Hispanic mutation and haplotype. Hum. Mutat. 27, 31–43 (2006).

35. Yuan, X. et al. Biomarkers of mature neuronal differentiation and related diseases. Future Sci. OA 10, 2410146 (2024).

36. Hill, B. G. et al. Integration of cellular bioenergetics with mitochondrial quality control and autophagy. Biol. Chem. 393, 1485–1512 (2012).

37. Marchetti, P., Fovez, Q., Germain, N., Khamari, R. & Kluza, J. Mitochondrial spare respiratory capacity: Mechanisms, regulation, and significance in non-transformed and cancer cells. FASEB J. 34, 13106– 13124 (2020).

38. Costanzo, M., et al. Proteomics Reveals that Methylmalonyl-CoA Mutase Modulates Cell Architecture and Increases Susceptibility to Stress. Int. J. Mol. Sci. 21, 4998 (2020).

39. Granath-Panelo, M. & Kajimura, S. Mitochondrial heterogeneity and adaptations to cellular needs. Nat. Cell Biol. 26, 674–686 (2024).

40. Sieber, M. H. & Spradling, A. C. The role of metabolic states in development and disease. Curr. Opin. Genet. Dev. 45, 58–68 (2017).

41. Oh, S. L., Cheng, L. Y., J Zhou, J. F., Henke, W. & Hagen, T. Galactose 1 -phosphate accumulates to high levels in galactose-treated cells due to low GALT activity and absence of product inhibition of GALK. J. Inherit. Metab. Dis. 43, 529–539 (2020).

42. Skolik, R. A., Solocinski, J., Konkle, M. E., Chakraborty, N. & Menze, M. A. Global changes to HepG2 cell metabolism in response to galactose treatment. Am. J. Physiol.-Cell Physiol. 320, C778–C793 (2021).

43. Protasoni, M. & Taanman, J.-W. Remodelling of the Mitochondrial Bioenergetic Pathways in Human Cultured Fibroblasts with Carbohydrates. Biology 12, 1002 (2023).

44. Gray, L. R., Tompkins, S. C. & Taylor, E. B. Regulation of pyruvate metabolism and human disease. Cell. Mol. Life Sci. 71, 2577–2604 (2014).

45. Zhdanov, A. V., Waters, A. H. C., Golubeva, A. V., Dmitriev, R. I. & Papkovsky, D. B. Availability of the key metabolic substrates dictates the respiratory response of cancer cells to the mitochondrial uncoupling. Biochim. Biophys. Acta BBA - Bioenerg. 1837, 51–62 (2014).

46. Morath, M. A. et al. Neurodegeneration and chronic renal failure in methylmalonic aciduria—A pathophysiological approach. J. Inherit. Metab. Dis. 31, 35–43 (2008).

47. Longo, N., Sass, J. O., Jurecka, A. & Vockley, J. Biomarkers for drug development in propionic and methylmalonic acidemias. J. Inherit. Metab. Dis. 45, 132–143 (2022).

48. Ribas, G. S., Lopes, F. F., Deon, M. & Vargas, C. R. Hyperammonemia in Inherited Metabolic Diseases. Cell. Mol. Neurobiol. 42, 2593–2610 (2022).

49. Halperin, M. L., Schiller, C. M. & Fritz, I. B. The inhibition by methylmalonic acid of malate transport by the dicarboxylate carrier in rat liver mitochondria. J. Clin. Invest. 50, 2276–2282 (1971).

50. Cheema-Dhadli, S., Leznoff, C. C. & Halperin, M. L. Effect of 2-Methylcitrate on Citrate Metabolism: Implications for the Management of Patients with Propionic acidemia and Methylmalonic aciduria. Pediatr. Res. 9, 905–908 (1975).

51. Mirandola, S. R. et al. Methylmalonate inhibits succinate-supported oxygen consumption by interfering with mitochondrial succinate uptake. J. Inherit. Metab. Dis. 31, 44–54 (2008).

52. Okun, J. G. et al. Neurodegeneration in Methylmalonic Aciduria Involves Inhibition of Complex II and the Tricarboxylic Acid Cycle, and Synergistically Acting Excitotoxicity. J. Biol. Chem. 277, 14674–14680 (2002).

53. Brusque, A. M. et al. Inhibition of the mitochondrial respiratory chain complex activities in rat cerebral cortex by methylmalonic acid. Neurochem. Int. 40, 593–601 (2002).

54. Pettenuzzo, L. F. et al. Differential inhibitory effects of methylmalonic acid on respiratory chain complex activities in rat tissues. Int. J. Dev. Neurosci. 24, 45–52 (2006).

55. Costa, R. T., Santos, M. B., Alberto-Silva, C., Carrettiero, D. C. & Ribeiro, C. A. J. Methylmalonic Acid Impairs Cell Respiration and Glutamate Uptake in C6 Rat Glioma Cells: Implications for Methylmalonic Acidemia. Cell. Mol. Neurobiol. 43, 1163–1180 (2023).

56. Proctor, E. C., et al. The Effect of Methylmalonic Acid Treatment on Human Neuronal Cell Coenzyme Q10 Status and Mitochondrial Function. Int. J. Mol. Sci. 21, 9137 (2020).

57. Kölker, S. et al. Methylmalonic Acid, a Biochemical Hallmark of Methylmalonic Acidurias but No Inhibitor of Mitochondrial Respiratory Chain. J. Biol. Chem. 278, 47388–47393 (2003).

58. Vihinen, M. Functional effects of protein variants. Biochimie 180, 104–120 (2021).

59. Rossi, A. et al. Genetic compensation induced by deleterious mutations but not gene knockdowns. Nature 524, 230–233 (2015).

60. El-Brolosy, M. A. & Stainier, D. Y. R. Genetic compensation: A phenomenon in search of mechanisms. PLOS Genet. 13, e1006780 (2017).

61. El-Brolosy, M. A. et al. Genetic compensation triggered by mutant mRNA degradation. Nature 568, 193– 197 (2019).

62. El-Brolosy, M. A. et al. Mechanisms linking cytoplasmic decay of translation-defective mRNA to transcriptional adaptation. Science 391, eaea1272 (2026).

63. Rambout, X. & Maquat, L. E. Keeping cells fit. Science 391, 657–658 (2026).

64. Falcucci, L. et al. Transcriptional adaptation upregulates utrophin in Duchenne muscular dystrophy. Nature 639, 493–502 (2025).

65. Dietz, H. C. et al. Four Novel FBN1 Mutations: Significance for Mutant Transcript Level and EGF-like Domain Calcium Binding in the Pathogenesis of Marfan Syndrome. Genomics 17, 468–475 (1993).

66. Kelly, M. A. et al. Adaptation and validation of the ACMG/AMP variant classification framework for MYH7 - associated inherited cardiomyopathies: recommendations by ClinGen’s Inherited Cardiomyopathy Expert Panel. Genet. Med. 20, 351–359 (2018).

67. Keyfi, F. et al. Mutation analysis of genes related to methylmalonic acidemia: identification of eight novel mutations. Mol. Biol. Rep. 46, 271–285 (2019).

68. Yu, Y. et al. Different mutations in the *MMUT* gene are associated with the effect of vitamin B12 in a cohort of 266 Chinese patients with mut-type methylmalonic acidemia: A retrospective study. Mol. Genet. Genomic Med. 9, e1822 (2021).

69. Deodato, F., Boenzi, S., Santorelli, F. M. & Dionisi-Vici, C. Methylmalonic and propionic aciduria. Am. J. Med. Genet. C Semin. Med. Genet. 142C, 104–112 (2006).

70. Hörster, F. et al. Long-Term Outcome in Methylmalonic Acidurias Is Influenced by the Underlying Defect (mut0, mut−, cblA, cblB). Pediatr. Res. 62, 225–230 (2007).

71. Hörster, F. et al. Prediction of outcome in isolated methylmalonic acidurias: combined use of clinical and biochemical parameters. J. Inherit. Metab. Dis. 32, 630–639 (2009).

72. Froese, D. S. et al. Structures of the Human GTPase MMAA and Vitamin B12-dependent Methylmalonyl-CoA Mutase and Insight into Their Complex Formation. J. Biol. Chem. 285, 38204–38213 (2010).

73. Mascarenhas, R. et al. Architecture of the human G-protein-methylmalonyl-CoA mutase nanoassembly for B12 delivery and repair. Nat. Commun. 14, 4332 (2023).

74. Larsen-Ledet, S., Panfilova, A. & Stein, A. Variant effects on protein-protein interactions: methods, models and diseases. Preprint at 10.48550/ARXIV.2507.17446 (2025).

75. Zhong, Q. et al. Edgetic perturbation models of human inherited disorders. Mol. Syst. Biol. 5, MSB200980 (2009).

76. Sahni, N. et al. Widespread Macromolecular Interaction Perturbations in Human Genetic Disorders. Cell 161, 647–660 (2015).

77. Sahni, N. et al. Edgotype: a fundamental link between genotype and phenotype. Curr. Opin. Genet. Dev. 23, 649–657 (2013).

78. Plessl, T. et al. Protein destabilization and loss of protein-protein interaction are fundamental mechanisms in *cblA* -type methylmalonic aciduria. Hum. Mutat. 38, 988–1001 (2017).

79. Nunnari, J. & Suomalainen, A. Mitochondria: In Sickness and in Health. Cell 148, 1145–1159 (2012).

80. Monzel, A. S., Enríquez, J. A. & Picard, M. Multifaceted mitochondria: moving mitochondrial science beyond function and dysfunction. Nat. Metab. 5, 546–562 (2023).

81. Mailloux, R. J., Treberg, J., Grayson, C., Agellon, L. B. & Sies, H. Mitochondrial function and phenotype are defined by bioenergetics. Nat. Metab. 5, 1641–1641 (2023).

82. Zong, Y. et al. Mitochondrial dysfunction: mechanisms and advances in therapy. Signal Transduct. Target. Ther. 9, 124 (2024).

83. Frega, M. et al. Rapid Neuronal Differentiation of Induced Pluripotent Stem Cells for Measuring Network Activity on Micro-electrode Arrays. J. Vis. Exp. 54900 (2017) doi:10.3791/54900.

84. Shan, X. et al. Fully defined NGN2 neuron protocol reveals diverse signatures of neuronal maturation. *Cell Rep*. Methods 4, 100858 (2024).

85. Han, L.-S. et al. Clinical features and MUT gene mutation spectrum in Chinese patients with isolated methylmalonic acidemia: identification of ten novel allelic variants. World J. Pediatr. 11, 358–365 (2015).

86. Fernández-Lainez, C. et al. Isolated methylmalonic acidemia in Mexico: Genotypic spectrum, report of two novel MMUT variants and a possible synergistic heterozygosity effect. Mol. Genet. Metab. Rep. 41, 101155 (2024).

