## Supporting information for "Disturbed mitochondrial energy production in methylmalonic aciduria is cell-type and variant-dependent"

#### Table S1. Antibody list for immunocytochemistry

| Target | Species | Dilution | Incubation | Manufacturer | Catalogue number |
|---|---|---|---|---|---|
| Synaptophysin | rabbit | 1:2000 | ON, 4°C | abcam | ab32127 |
| MAP2 | chicken | 1:5000 | ON, 4°C | abcam | ab5992 |
| NeuN | mouse | 1:500 | ON, 4°C | abcam | ab104224 |
| Secondary antibodies | | | | | |
| Alexa Fluor™ 647 donkey-anti rabbit IgG (H+L) | donkey | 1:100 | 1h, RT | Thermo Fisher Scientific | A-31573 |
| Alexa Fluor™ 488 donkey-anti chicken IgG (H+L) | donkey | 1:100 | 1h, RT | Thermo Fisher Scientific | A78948 |
| Alexa Fluor™ 568 donkey-anti mouse IgG (H+L) | donkey | 1:100 | 1h, RT | Thermo Fisher Scientific | A10037 |

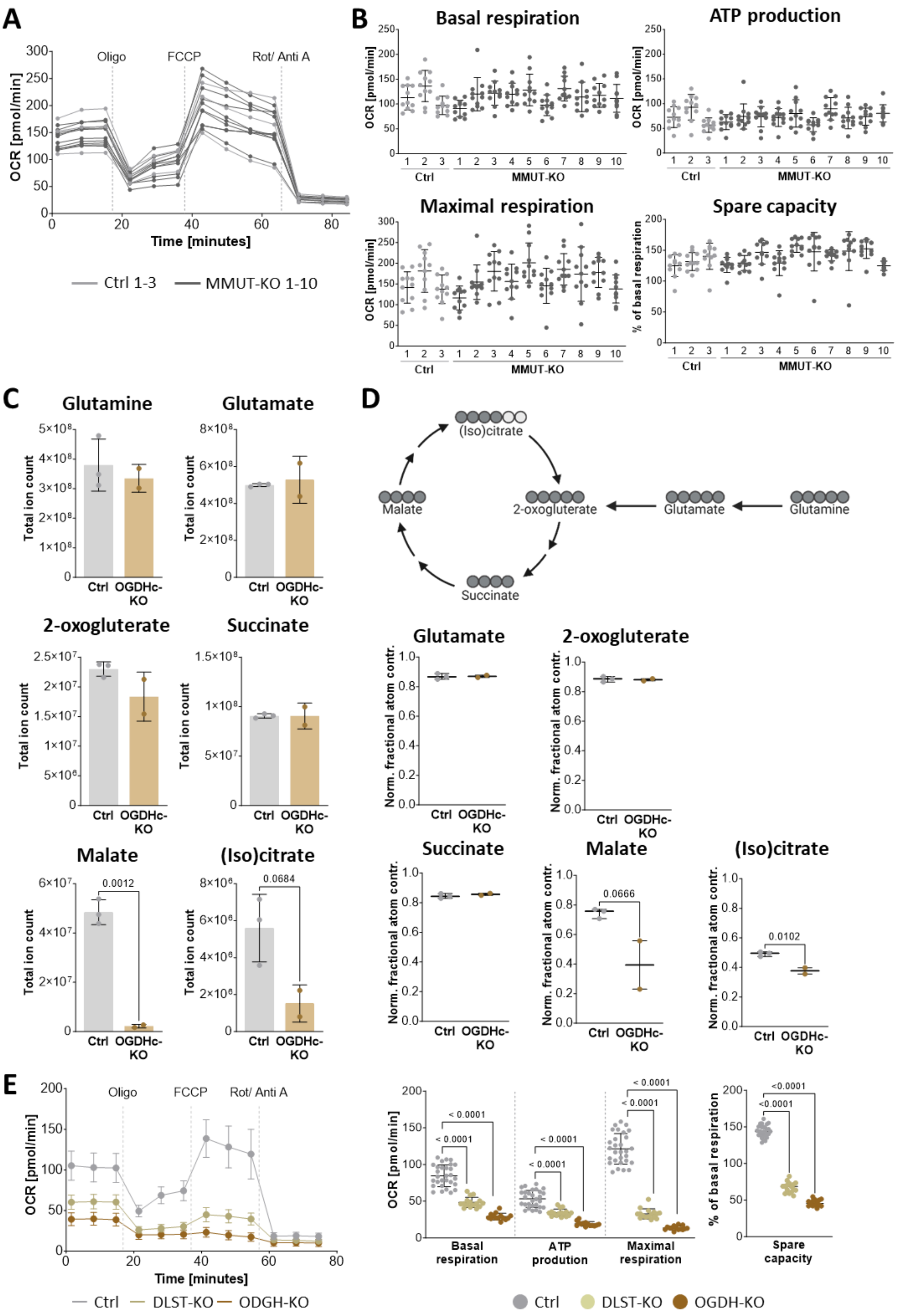

**Figure S1. Decreased mitochondrial energy production in 293T cells lacking OGDC subunits, but inconclusive results in MMUT-KOs.** (A) Normalized OCR profile from one representative experiment in 293T cells (n=3 for Ctrl and n=10 for MMUT-KO cells). Data is shown as mean ± SD.

Technical replicates were pooled. (B) Normalized parameters from eleven experiments in Ctrl (n=3) and MMUT-KO (n=10) cells. Data is shown as mean ± SD. One dot represents the mean of 2-3 technical replicates. (C) Pool levels of selected metabolites of the TCA cycle and related metabolites in Ctrl cells (n=3) and OGDHc-KO cells (n=2). Data is shown as mean ± SD. One dot represents one biological replicate based on the mean of two technical replicates. Unpaired t-test were used to compare OGDHc-KO to Ctrl cells (D) Top: Schematic for labelled TCA cycle metabolites after treatment with $^{13}$C-glutamine for 4 hours. Dark grey circles represent labelled C atoms. Bottom: Normalized fractional atom contribution of $^{13}$C-glutamine to TCA cycle metabolites after treatment with $^{13}$C-glutamine for 4 hours. Data is shown as mean ± SD, mean of biological replicates is based on two technical replicates. Unpaired t-test were used to compare OGDHc-KO to Ctrl cells. (E) Normalized OCR profile and parameters from one representative experiment in Ctrl cells (n=2) and DLST- and OGDH-KO cells (each n=1). Data is shown as mean ± SD. Dots represent technical replicates. For OCR profiles, technical replicates (n=14-16) and biological replicates (for Ctrl cells) were combined. Multiple unpaired t-tests were used to compare OGDHc-KO to Ctrl cells.

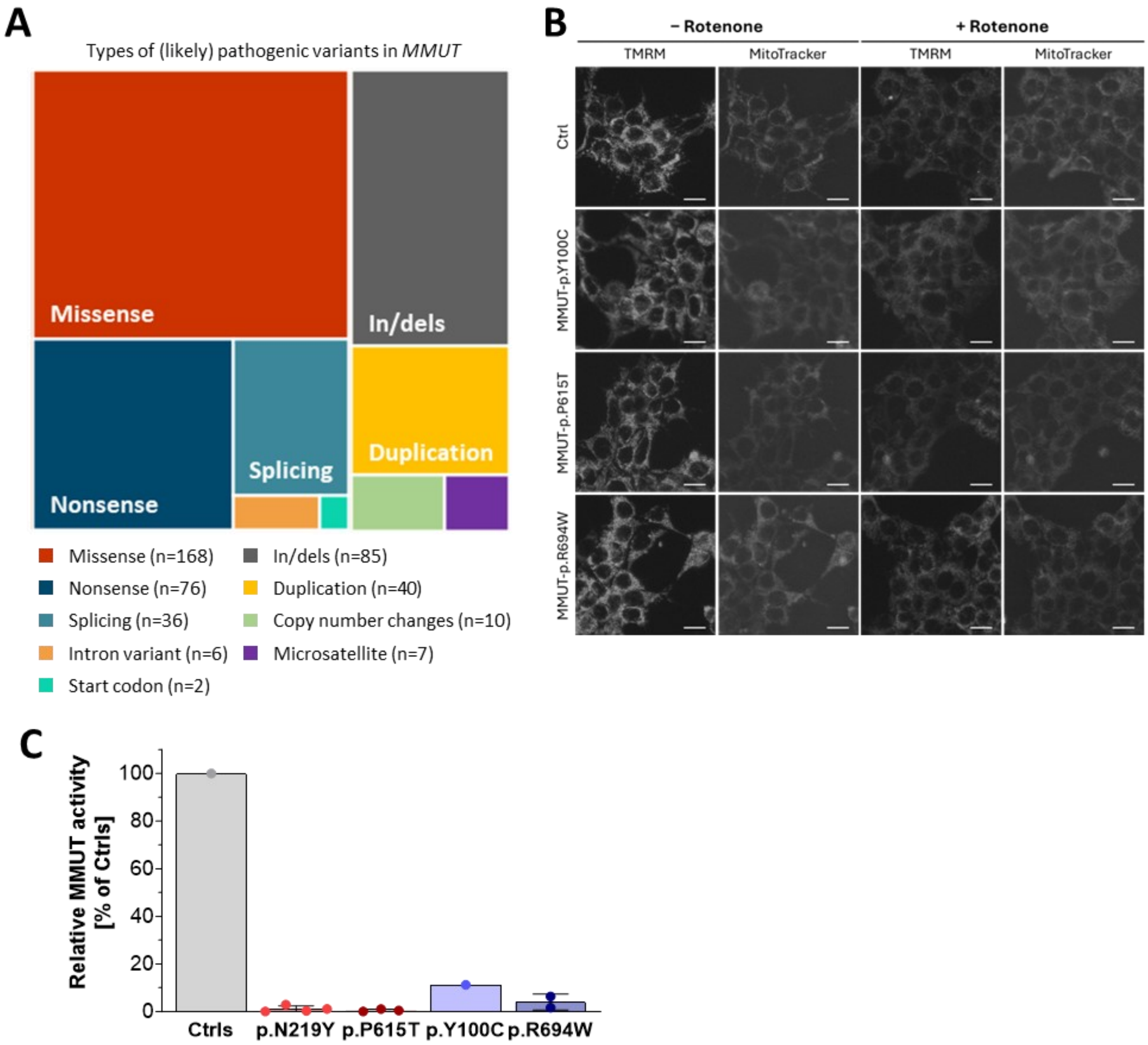

**Figure S2. Decreased MMUT protein levels and activity in 293T cells and fibroblasts wih MMUT missense variants.** (A) Proportions of molecular consequences of (likely) pathogenic single nucleotide changes in the *MMUT* gene as listed on ClinVar (June 2026). (B) Representative images of TMRM and MitoTracker Green measurements in the absence and presence of rotenone (1 µM) from one clone with MMUT-p.Y100C, MMUT-p.P615T and MMUT-p.R694W in 293T cells. Scale bar is 20 µm. (C) Relative MMUT enzymatic activity in fibroblasts compared to control (1'270 pmol/min/mg protein). Data is shown as mean ± SD and taken from previous studies[1]. One dot represents a biological replicate based on two technical replicates.

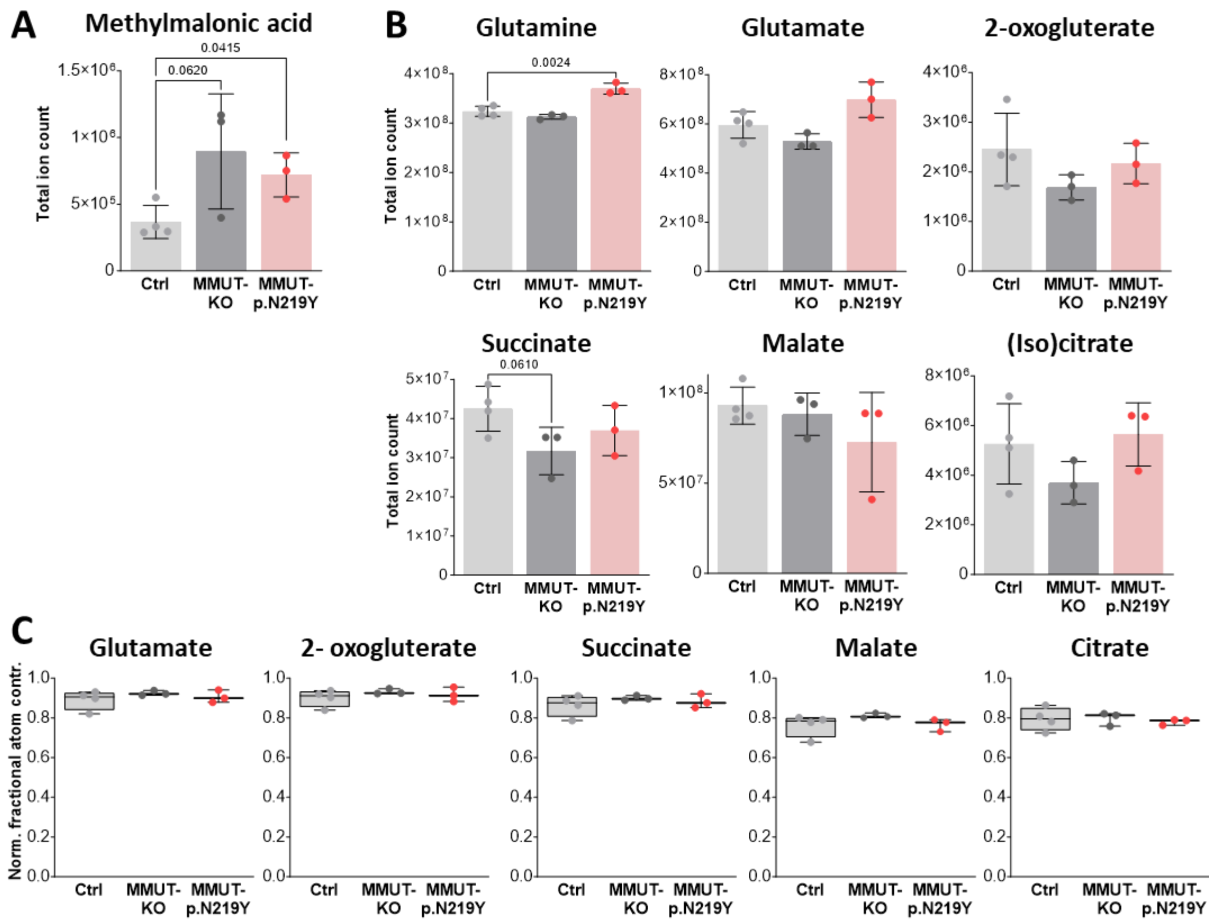

**Figure S3. Altered glutamine anaplerosis in MMUT-KO and MMUT-p.N219Y 293T cells.** (A) Total ions counts for methylmalonic acid in Ctrl (n=4; Ctrl 1-4), MMUT-KO (n=3; MMUT-KO1, KO2 and KO6) and MMUT-p.N219Y (n=3) 293T cells. Data is shown as mean ± SD. Dots represent biological replicates, based on the mean of two technical replicates. p-values were calculated with unpaired t-tests. (B) Total ion counts for TCA cycle metabolites acid in Ctrl, MMUT-KO and MMUT-p.N219Y 293T cells. Data is shown as mean ± SD. Dots represent biological replicates, based on the mean of two technical replicates. p-values were calculated with unpaired t-tests. (C) Normalized fractional atom contribution of $^{13}$C-glutamine to TCA cycle metabolites after treatment with $^{13}$C-glutamine for 4 hours. in Ctrl, MMUT-KO and MMUT-p.N219Y 293T cells. Data is shown as mean ± SD, mean of biological replicates is based on two technical replicates.

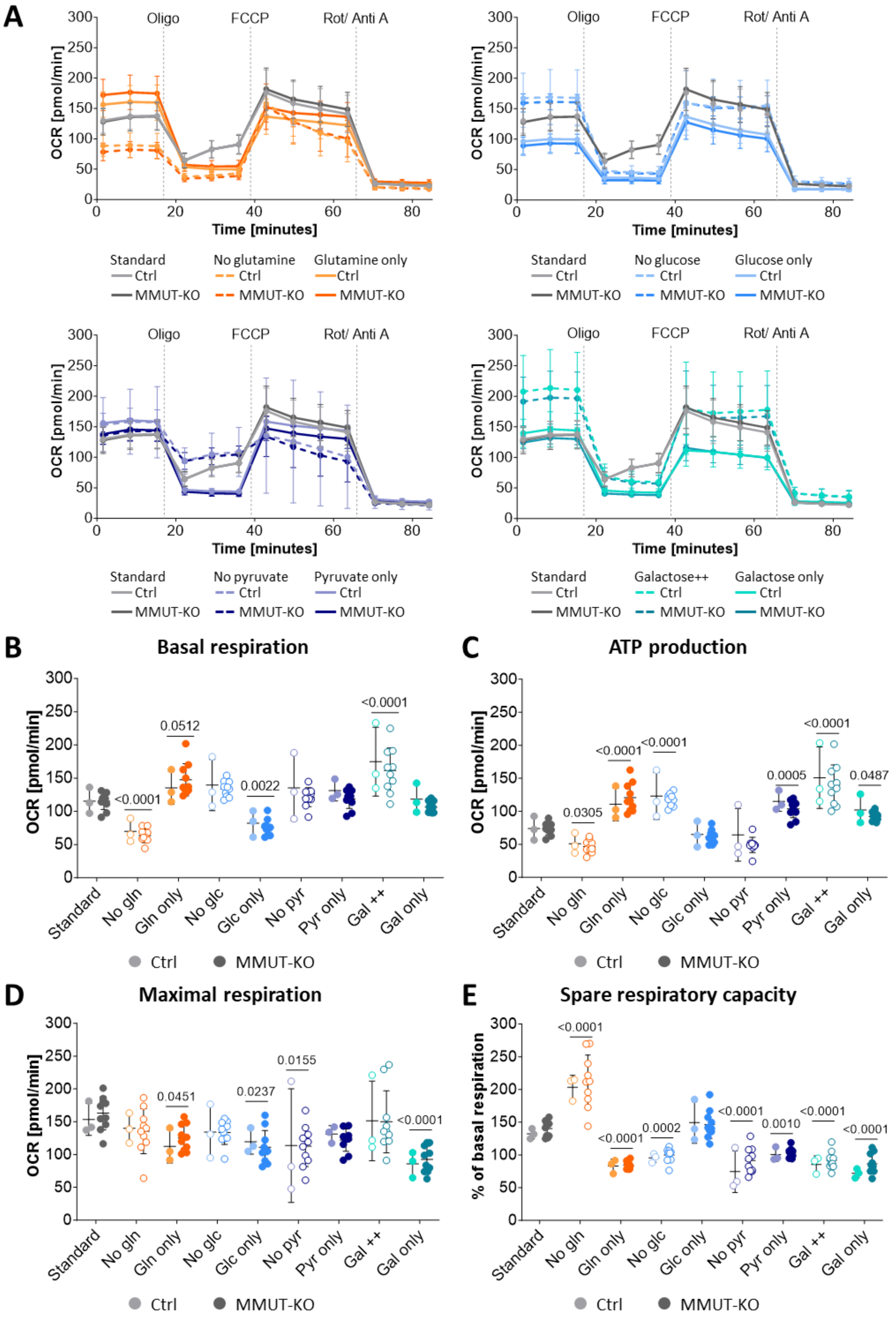

**Figure S4. Examination of TCA cycle fuel sources on mitochondrial energy production in MMUT-KO 293T cells.** (A) Normalized OCR profiles in Ctrl (n=3) and MMUT-KO (n=19) 293T cells under standard conditions (2.0 mM glutamine, 2.0 g/l glucose, 1.0 mM pyruvate) and stress conditions (0.25 mM glutamine in glutamine only; 0.25 g/l glucose for glucose only; 0.125 mM pyruvate for pyruvate only; 2.0 g/l galactose for galactose++ and 0.25 g/l for galactose only. Colors refer to TCA cycle fuel sources depicted in Figure 4A, with light shades for Ctrl and darker shades for MMUT-KO clones. Biological replicates were pooled for OCR profiles, and profiles for standard conditions are pooled from eleven experiments. (B-E) Normalized parameters under standard and stress conditions. Each dot represents one biological replicate based on the mean of 2-4 technical replicates, or the mean of eleven experimental replicates with 2-4 technical replicates for standard conditions. A two-way ANOVA with Dunnett's multiple comparisons test was used to compare the mean of MMUT-KO and Ctrl cells in each media condition to the mean under standard conditions, reporting the significant adjusted p-values ($p < 0.05$) groups and conditions. Abbreviations: Gln: glutamine; Glc: glucose; Pyr: pyruvate; Gal: galactose.

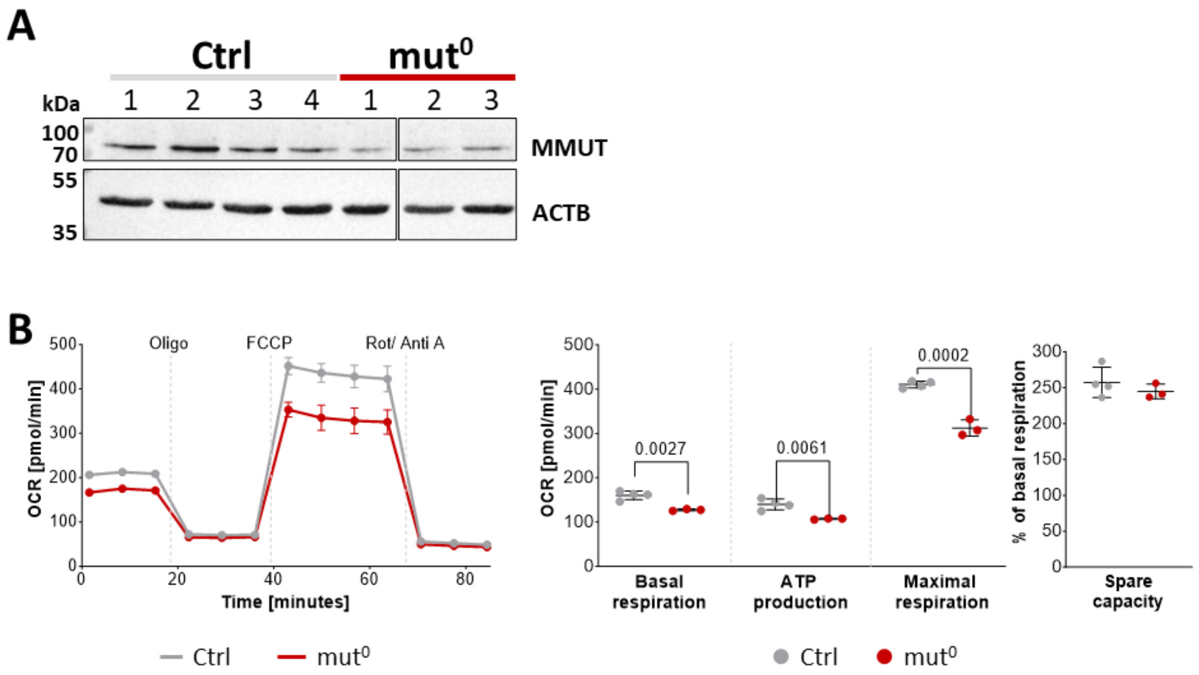

**Figure S5. Urine derived kidney cells show decreased mitochondrial energy production.** (A) Western Blot against MMUT and quantification in Ctrl (n=4) and MMUT-deficient urine-derived kidney cells. ACTB was used as loading control. Data is shown as mean ± SD. An unpaired t-test was used to compare groups. (B) Normalized OCR profiles and individual parameters of Ctrl (n=4) and MMUT-deficient (n=3) urine-derived kidney cells. Data is shown as mean ± SD. Each dot represents a biological replicate, based on the mean of 11-12 technical replicates from two experimental replicates. Biological replicates were pooled for OCR profiles. An unpaired t-test was applied. This data replicates previously published results[2,3].

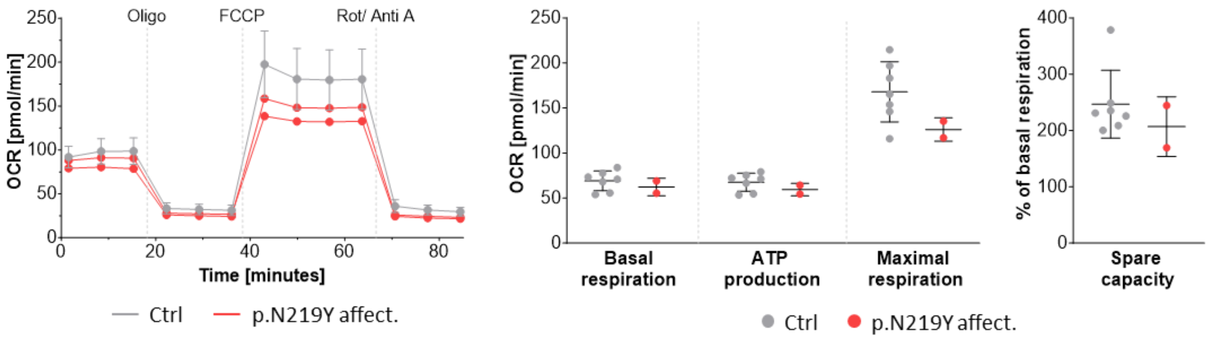

**Figure S6.** Normalized OCR profiles and individual parameters in Ctrl (n=7) and MMUT-p.N219Y (n=2) fibroblasts. Data is shown as mean ± SD. Each dot represents a biological replicate, based on the mean of 3-8 technical replicates from two experimental replicates. Biological replicates were pooled for OCR profiles. The MMUT-deficient cell lines were used for reprogramming into iPSCs[4].

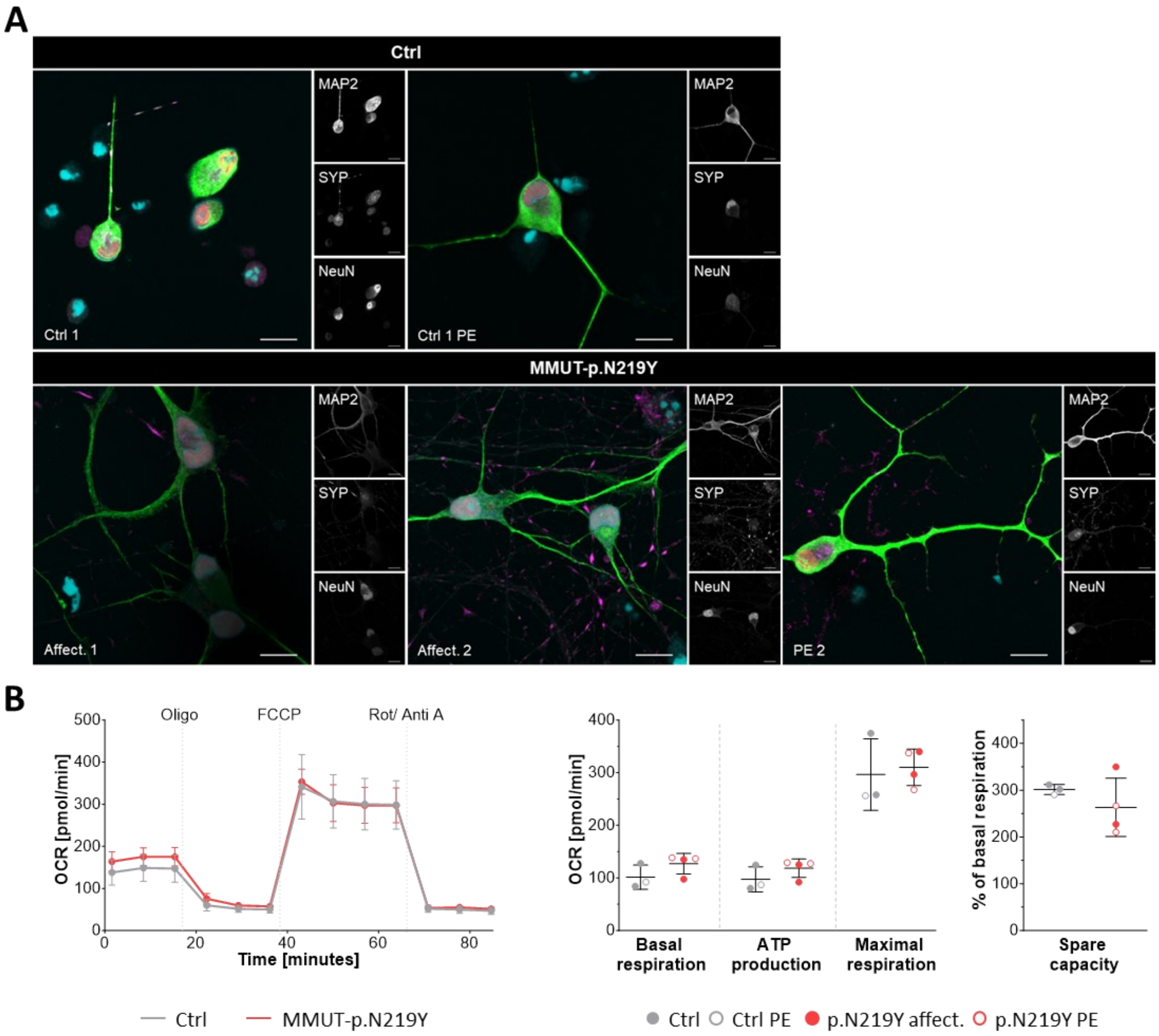

**Figure S7. MMUT-p.N219Y does not influence energy production in neurons at day 21 of differentiation.** (A) Representative immunocytochemistry images of neurons at day 14 of differentiation of Ctrl (top) and MMUT-p.N219Y (bottom). Shown are microtuble associated protein 2 (MAP2, green), Synaptophysin (SYP, magenta), NeuN (red) and nuclei stained with Hoechst (cyan). Scale bar is 10 µm. (B) OCR profiles and individual parameters of Ctrl (n=3) and MMUT-p.N219Y clones (n=4) in neurons at day 21 of differentiation. Data is shown as mean ± SD. Each dot represents a biological replicate, based on the mean of 2-11 technical replicates from three experimental replicates. Biological replicates were pooled for OCR profiles. Abbreviations: affect. = Affected cell line; PE = prime edited.

#### Figure 1B

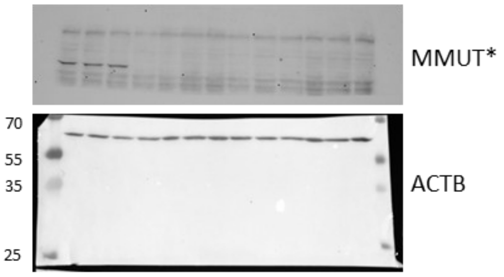

*no overlay with ladder available

#### Figure 2B

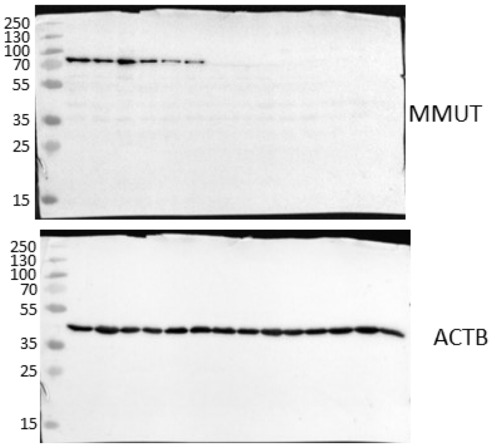

ACTB developed after stripping of membrane

#### Figure 3B/ S2B

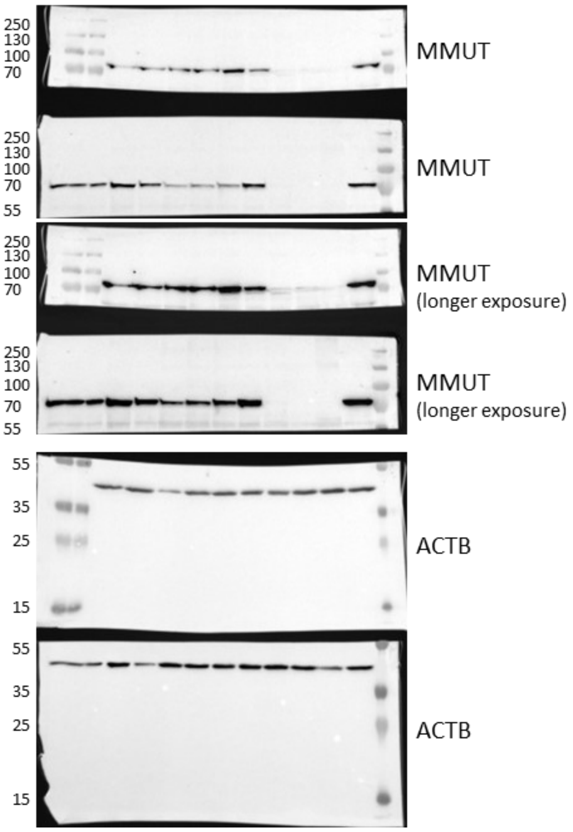

#### Figure 3G/ S2D

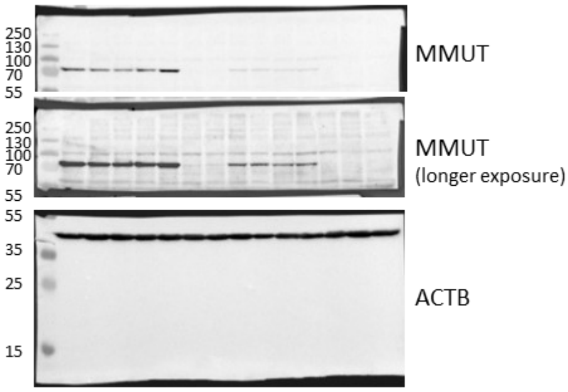

#### Figure S5

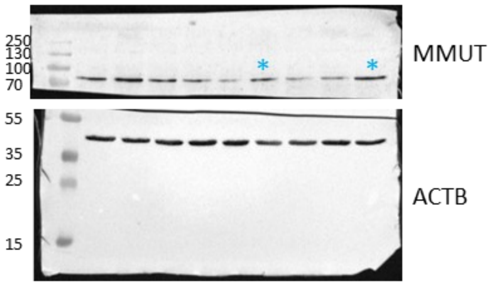

*Cell line got contaminated with control cell line and was excluded from further experiments

#### Figure 5B

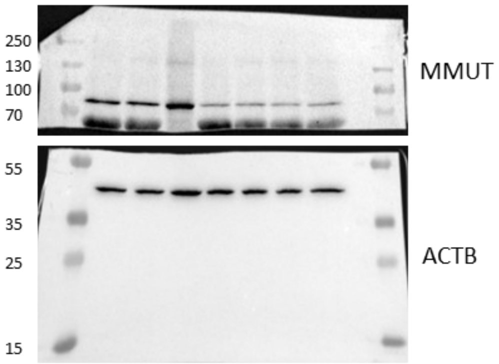

**Figure S8. Uncropped membranes for all Western blot experiments performed in this study.**

#### Supplementary Methods

##### Generation of Knockout cells lines and prime editing

293T cells with knockouts of DLST, OGDH as well MMUT-KO1 and KO2 were previously generated[1] and further characterized in this publication. CRISPR-Cas9 editing with homology directed repair (HDR) was performed in 293T cells to generate MMUT-KO3-10 and MMUT-p.Y100C clones. Cas9 protein (TrueCut™ Cas9 v2, Thermo Fisher scientific, A36498) and sgRNAs were provided as ribonucleoprotein complex and templates for HDR as single-stranded oligodeoxynucleotides (ssODNs). sgRNAs were synthesized by in vitro transcription using the GeneArt™ Precision gRNA Synthesis Kit (Invitrogen, A29377). Sequences of the sgRNAs and ssODNs can be found in the table below.

| | **Sequence sgRNA 5'-3'** |
|---|---|
| sgRNA to generate MMUT-KO3, MMUT-KO4 and MMUT-p.Y100C | GCGGATGGTCCAGGGCC TAA AGG |
| sgRNA to generate MMUT-KO5 and MMUT-KO6 | GTACCTAAAGAGAAG CTT ACT GG |
| sgRNA to generate MMUT-KO7 - MMUT-KO10 | AAGAATATCTGGCCGTC CAA GGG |
| ssODN for MMUT-p.Y100C | CATTCACACGTGGACCATATCCTACCATGTGTACTTTTAGGCCC TGGACCATCCGCCAGTATGCTGGTTTTAGTACTGTGGAAGAAA GCAATAAGTTCTATAAGGACAACATTAAGGGTGAGATTTT |

293T cells were transfected by electroporation (1150 V for 20 ms with 2 pulses) using a Neon transfection system (Thermo Fisher Scientific) containing 150'000 cells, 243.6 ng sgRNA, 312.5 ng Cas9 (Invitrogen, A36498) and 20 pmol ssODN following the manufacturer's instructions. After 48 h, cells were collected with trypsin and diluted to 1 cell per 100 µl for single cell seeding a 96-well plate. Positive clones were confirmed by Sanger sequencing (for sequences see table below)

| | HGVS Description (cDNA) NC_000006.12(NM_000255.3) | HGVS Description (protein) NC_000006.12(NP_000246.2) |
|---|---|---|
| **KOs in 293T cells** | | |
| MMUT-KO1 | c.656_668del and c.656dup | p.N219Rfs*17 and p.N219Kfs*2 |
| MMUT-KO2 | c.657dup and c.654_656delinsGTAAT | p.N219* and p.D220* |
| MMUT-KO3 | c.309dup | p.(Pro104Alafs*11) |
| MMUT-KO4 | c.309dup | p.(Pro104Alafs*11) |
| MMUT-KO5 | c.638_639insA; c.639dup | p.(Thr214Tyrfs*7) |
| MMUT-KO6 | c.639dup | p.(Thr214Tyrfs*7) |
| MMUT-KO7 | c.2078del | p.(Gly693Aspfs*12) |
| MMUT-KO8 | c.2064_2079del | p.(Glu688Aspfs*12) |
| MMUT-KO9 | c.2078_2086del | p.(Gly693_Pro695del) |
| MMUT-KO10 | c.2078_2095del | p.(Gly693_Leu698del) |
| **KOs in fibroblasts** | | |
| MMUT-KO1 MMUT-KO2 | c.88C>T | p.Q30* |
| MMUT-KO3 MMUT-KO4 | c.1207C>T | p.Arg403* |
| MMUT-KO5 MMUT-KO6 | c.1531C>T | p.Arg511* |
| MMUT-KO7 MMUT-KO8 | c.2179C>T | p.Arg727* |
| **Missense variants in 293T cells and fibroblasts** | | |
| MMUT-p.Y100C | c.299A>G | p.Tyr100Cys |
| MMUT-p.N219Y | c.655A>T | p.Asn219Tyr |
| MMUT-p.P615T | c.1843C>A | p.Pro615Thr |
| MMUT-p.R694W | c.2080C>T | p.Arg694Trp |

Prime-editing was performed in 293T cells and iPSCs to introduce the variants MMUT-p.N219Y, MMUT-p.P615T and/or MMUT-p.R694W, using the PE3 system[5]. 293T cells were transfected with Lipofectamine 3000, using 1’500 ng PEmax plasmid (pCMV-PEmax-P2A-BSD was a gift from David Liu; Addgene, #174821[6]), 500 ng epegRNA (in pU6-tevopreq1-GG-acceptor; a gift from David Liu; Addgene, #174038[7]) and 166 ng nicking sgRNA (in pAT9658-sgRNA-mCherry; a gift from Ervin Welker; Addgene, #162987[8]). iPSCs were transfected by electroporation (950V/ 40ms/ 1 pulse) with 375 ng pCMV-PEmax (a gift from David Liu; Addgene, #174820[6]), 125 ng epegRNA and 41.5 ng nicking guide sgRNA. Sequences of the epegRNA and nicking guide RNA can be found in the table below. Cell populations were analyzed 72 h post-transfection using Sanger sequencing or kept in culture for blasticidin (InvivoGen, ant-bl-1) selection for 11 days and then analyzed by Sanger sequencing.

| | |
|---|---|
| **epegRNA protospacer** | |
| MMUT-p.N219Y | TTCCTTTAGTATATCATTT |
| MMUT-p.P615T | TTTTTGCTACAAGAAGACG |
| MMUT-p.R694W | AGAACTTAACTCCCTTGGA |
| **nicking guide RNA** | |
| MMUT-p.N219Y -40 | TAGTAACTGGAGAAGAACA |
| MMUT-p.P615T +41 | GACAAGATGGCCATGACAG |
| MMUT-p.R694W +3 | AGAATATCTGGCCATCCAA |

**Western Blotting**

Cells were collected by trypsination, centrifuged, and washed once with DPBS. Cell pellets were either stored at -80°C or immediately lysed by resuspending in RIPA buffer (150 mM NaCl, 50 mM Trizma-Base (pH 8), 1% NP-40, 10% sodium deoxycholate and 1% Halt™ Protease & Phosphatase Inhibitor Cocktail (100X, Thermo Fisher Scientific, 784409)), incubated on ice for 30 min and centrifugated at 17’000*g* at 4°C for 5 min to remove cell debris. Protein concentration was determined using a BCA assay (Thermo Fisher Scientific; 23227). Protein lysates were diluted with lysis buffer and 4x Laemmli buffer (Bio-Rad, 161-0747) with a final concentration of 5% β-mercaptoethanol (Sigma, M6250) and incubated at 96°C for 5 min and then placed on ice. 20 µg of protein per sample was loaded on self-cast 12% SDS-gels and electrophoresed at 140 V until the dye front reached the end of the gel. Proteins were transferred onto a 0.45 µm nitrocellulose membrane (Cytiva, 10600007) using the Trans-Blot®Turbo™ Transfer System (Bio-Rad) and 1x transfer buffer (10x stock: 25 mM TrisBase, 192 mM glycine) with 20% methanol and 0.2% SDS. Successful transfer was confirmed by staining with Ponceau S solution (Sigma-Aldrich, P7170-

1L). Membranes were blocked at ambient temperature for at least 60 min in 5% milk powder (w/v; Millipore, 70166-500 G) in 1x Tris-buffered saline (TBS-T; 20 mM TrisBase, 150 mM NaCl) with 0.2% Tween20 (v/v; Sigma, P1379-100ML). Incubation with primary antibodies (MMUT: abcam, ab67869, 1:500; ACTB: Sigma-Aldrich, A1978, 1:1000) was performed overnight at 4°C while shaking. After washing three times with TBS-T (5 min at room temperature while shaking), membranes were incubated with the secondary antibody (anti-mouse horseradish peroxidase: SantaCruz, sc516102-cm, 1:5000) for 1 hour at room temperature, followed by three washes with TBS-T. Chemiluminescent signals were developed using Clarity Max™ Western ECL Substrate (Bio-Rad, 1705062) or SuperSignal™ West Femto Maximum Sensitivity Substrate (Thermo Fisher Scientific, 34096) and visualized on a ChemiDoc™ XRS (Bio-Rad) with Image Lab (Bio-Rad, Version 6.1.0).

**Differentiation of iPSCs into neurons**

iPSCs were differentiated into glutaminergic excitatory neurons using NGN2-induced lentiviral programming, as previously described[9]. Briefly, iPSCs were seeded at a density of 350’000 to 430’000 cells in StemFlex medium, 24 h prior to lentiviral infection.

The lentiviral vectors used were FUW-M2-rtTA (reverse tetracycline-controlled transactivator, a gift from Rudolf Jaenisch; Addgene, #20342[10]) and tet-O-Ngn2-puro (doxycycline-inducible NGN2, a gift from Marius Wernig; Addgene, #52047[9]). All lentiviruses were produced in 293T cells using packaging plasmids pMDLg/pRRE (Addgene, #12251), pMD2.G (Addgene, #12259), and pRSV-Rev (Addgene, #12253) which were gifts from Didier Trono[11]. Briefly, 293T cells were co-transfected with the packaging plasmids and one lentivector. Virus-containing medium was collected at 24 and 48 h post-transfection and incubated overnight with PEG-6000 precipitation solution (Thermo Fisher Scientific, A17541.30) at 4°C. The precipitated virus was pelleted by centrifugation at 3’500g t 4°C for 1 h, rinsed with PBS and resuspended in PBS. The viral suspension was then aliquoted, snap-frozen, and stored at −80°C for long-term storage.

On Day 0 of differentiation (24 h post infection), 2.5 µg/ml doxycycline (Sigma-Aldrich, D9891) was added to induce NGN2 expression, and cells were selected on Days 1–2 in BrainPhys™ Neuronal Medium (Stemcell Technologies, 05790) supplemented with 2.5 µg/ml doxycycline and 2.5 µg/ml puromycin (Thermo Fisher Scientific, A11138-03). On Day 3, neurons were dissociated with Accutase (Gibco, A1110501), passed through a cell strainer (40 µm; Corning, 431750) and replated in 96-well plates for subsequent assays or in 8-well chamber slides (ibidi, 80841) in BrainPhys medium containing 2.5 µg/ml doxycycline and ROCK inhibitor (1:1000 from 10 mM stock, reconstituted in DMSO; STEMCELL Technologies, 72304). On Day 4, cells were maintained in BrainPhys medium with 2.5 µg/ml doxycycline. On Days 5–6, medium was supplemented with 2.5 µg/ml doxycycline, 10 ng/ml BDNF (Thermo Fisher Scientific, PHC7074), 10 ng/ml NT-3 (Thermo Fisher Scientific, PHC7036), and 5 µM AraC (Sigma-Aldrich, C1768). From Day 7 onward, cells were maintained in BrainPhys medium supplemented with 2.5 µg/ml doxycycline, 10 ng/ml BDNF, 10 ng/ml NT-3, and 20 ng/ml GDNF (Thermo Fisher Scientific, PHC7041), with half-medium changes performed every 2–3 days. rhLaminin-521 (Gibco, A29249) was added weekly to fresh prewarmed medium immediately before feeding to support neuronal attachment.

**Stable isotope tracing**

Stable isotope tracing in DLST-KO and OGDH-KO 293T cells was performed as previously published[1]. For MMUT-KO and MMUT-p.N219Y 293T cells, a total of

700'000 cells per well were seeded on poly-L-lysine (Sigma, P4707-50ML) coated 6-well plates in high glucose DMEM (Gibco, 31966-047) and allowed to attach overnight. The next day, cells were washed with DPBS and medium was exchanged to high glucose DMEM (Gibco, 11960) supplemented with 5% dialyzed FBS (Thermo Fisher Scientific, 26400044) and 4 mM $^{13}C_5$-glutamine (Merck, 605166). Cells were incubated for 4 h at 37°C, before washing with ice cold DPBS and incubation with 500 µl ice pre-cooled extraction solution (40:40:20 methanol: acetonitrile: $H_2O$ + 0.5% formic acid) for 2 min. Cells were collected using a scrapper and formic acid was quenched by adding 44 µl 15% $NH_4CO_3$ (w/v). Homogenates were incubated on ice for 20 min and centrifuged at 16'000g at 4°C for 10 min. Supernatants and pellets were snap-frozen in liquid nitrogen and stored at -80°C before analysis by liquid chromatography-mass spectrometry (LC-MS, see method section 2.11) and protein quantification, respectively. For protein quantification, pellets were resuspended in 200 µl of 200 mM NaOH, incubated at 95°C for 20 min and centrifuged at 2'000g for 10 min. A BCA assay was performed, using 10 µl of solubilized proteins, following manufacturer's instructions.

**Metabolite measurement by liquid chromatography-mass spectrometry**

Metabolites were measured on an Orbitrap Astral Mass Spectrometry (Thermo Fisher Scientific) which was coupled to hydrophilic interaction chromatography (HILIC) by electrospray ionization. During measurements, samples were stored at 4°C in the autosampler with 5 µl per sample injected. For HILIC separation, a XBridge BEH Amide column (130Å pore size, 2.5 µm, 2.1 mm X 150 mm; Waters™, 186006724) and a gradient of solvent A (95:5 water: acetonitrile with 10 mM ammonium acetate and 10 mM ammonium hydroxide, pH 9.45) and solvent B (acetonitrile) was used. The gradient was composed as followed: 0 min, 90% B; 3

min, 75% B; 8 min, 70% B; 10 min, 50% B; 13 min, 25% B; 16 min, 0% B; 22 min, 90% B; 27 min, 90% B. Flow rate of this 27-minutes method was 150 µl per minute. Measurements were acquired in negative ion mode with data-dependent acquisition and dynamic exclusion after one occurrence of 4 seconds. Full scans were measured in a window of m/z 70 to 1000 in the Orbitrap analyzer at 120'000 resolution, an AGC of 1e6 and ITmsx of 100 ms. Measurements of MS/MS spectra in the Astral analyzer were performed in a window of m/z 40-1000 with an AGC of 1e3 and ITmsx of 3 ms. The isolation window for precursors was 2 m/z and the HCD collision energy 30%. Time between master scans was 0.3 seconds.

Peak picking and analysis of metabolomics data was performed with the ElMaven software (https://github.com/ElucidataInc/ElMaven). Data was corrected for natural abundance of $^{13}C$ with Accucor (https://github.com/XiaoyangSu/AccuCor). Total ion counts describe the sum of ion counts detected for all isotopologues of the respective metabolite. Fractional contribution describes the atom contribution of the tracer to the metabolite pool of interest. The calculation was based on the definition in[12] and is described as:

$$FCatom = \frac{\sum_{i=0}^{N}(M+i) * i}{N * \sum_{i=0}^{N}(M+i)}$$

with $FC_{atom}$ = fractional contribution of the labelled atom, M+i abundance of isotopologues containing i labelled atoms, i = number of labelled atoms in that isotopologue and N = maximum number of tracer-relevant atoms in the metabolite. To correct for incomplete tracer enrichment in the tracer metabolite pool, the fractional contribution of the metabolite of interest was normalized to the fractional atom contribution of the tracer to the tracer metabolite pool and is defined as:

$$normalized\ FCatom = \frac{FCatom\ of\ metabolite}{FCatom\ of\ the\ tracer}$$

#### Analysis pipeline Harmony
#### 293T cells

| Input Image | Input |
|---|---|
| | **Flatfield Correction :** Basic<br>Brightfield Correction<br>**Stack Processing :** Individual Planes<br>**Min. Global Binning :** Dynamic |

| Find Nuclei | Input | Method | Output |
|---|---|---|---|
| | **Channel :** HOECHST 33342<br>**ROI :** None | **Method :** C<br>Common Threshold : 0.4<br>Area : > 30 μm²<br>Splitting Coefficient : 7.0<br>Individual Threshold : 0.4<br>Contrast : > 0.1 | Output Population : Nuclei |

| Find Cytoplasm | Input | Method | Output |
|---|---|---|---|
| | **Channel :** TMRE<br>**Nuclei :** Nuclei | **Method :** C<br>Common Threshold : 0.45<br>Individual Threshold : 0.15 | |

| Find Cytoplasm (2) | Input | Method | Output |
|---|---|---|---|
| | **Channel :** MitoTracker Green<br>**Nuclei :** Nuclei | **Method :** C<br>Common Threshold : 0.45<br>Individual Threshold : 0.15 | |

| Calculate Intensity Properties | Input | Method | Output |
|---|---|---|---|
| | **Channel :** TMRE<br>**Population :** Nuclei<br>**Region :** Cytoplasm | **Method :** Standard<br>Mean | Property Prefix : Intensity TMRM |

| Calculate Intensity Properties (2) | Input | Method | Output |
|---|---|---|---|
| | **Channel :** MitoTracker Green<br>**Population :** Nuclei<br>**Region :** Cytoplasm | **Method :** Standard<br>Mean | Property Prefix : Intensity MitoTracker Green |

| Define Results | Results |
|---|---|
| | **Method :** List of Outputs<br>**Population : Nuclei**<br>Number of Objects<br>Apply to All : Mean+StdDev<br>Intensity TMRM Mean : Mean+StdDev<br>Intensity MitoTracker Green Mean : Mean+StdDev<br><br>**Method :** Formula Output<br>Formula : a/b<br>Population Type : Objects<br>Variable a : Nuclei - Intensity TMRM Mean Mean<br>Variable b : Nuclei - Intensity MitoTracker Green Mean Mean<br>Output Name : TMRM/ Mito Tracker |

#### iPSC-derived neurons

| Input Image | Input | | |
|---|---|---|---|
| | **Flatfield Correction :** Basic<br>**Stack Processing :** Maximum Projection<br>**Min. Global Binning :** Dynamic | | |
| **Find Nuclei** | **Input** | **Method** | **Output** |
| | **Channel :** HOECHST 33342<br>**ROI :** None | **Method :** C<br>Common Threshold : 0.5<br>Area : > 30 µm²<br>Splitting Coefficient : 90<br>Individual Threshold : 0.8<br>Contrast : > 0.2 | Output Population : Nuclei |
| **Calculate Intensity Properties** | **Input** | **Method** | **Output** |
| | **Channel :** HOECHST 33342<br>**Population :** Nuclei<br>**Region :** Nucleus | **Method :** Standard<br>Mean | Property Prefix : Intensity Nucleus HOECHST 33342 |
| **Select Population** | **Input** | **Method** | **Output** |
| | **Population :** Nuclei | **Method :** Filter by Property<br>Intensity Nucleus HOECHST 33342 Mean : >= 5000 | Output Population : True Nuclei |
| **Find Cytoplasm** | **Input** | **Method** | **Output** |
| | **Channel :** TMRE<br>**Nuclei :** True Nuclei | **Method :** B<br>Common Threshold : 0.51<br>Individual Threshold : 0.3 | |
| **Calculate Intensity Properties (2)** | **Input** | **Method** | **Output** |
| | **Channel :** TMRE<br>**Population :** True Nuclei<br>**Region :** Cell | **Method :** Standard<br>Mean | Property Prefix : Intensity Cell TMRE |
| **Calculate Morphology Properties** | **Input** | **Method** | **Output** |
| | **Population :** True Nuclei<br>**Region :** Cell | **Method :** Standard<br>Area<br>Roundness | Property Prefix : Cell |

| Select Population (2) | Input | Method | Output |
|---|---|---|---|
| | **Population :** True Nuclei | **Method :** Filter by Property<br>Intensity Cell TMRE Mean : > 2900<br>Cell Area [µm²] : <= 700<br>Cell Area [µm²] : > 70<br>Boolean Operations : F1 and F2 and F3 | Output Population : Live cells |

| Select Cell Region | Input | Method | Output |
|---|---|---|---|
| | **Population :** Live cells | **Method :** Resize Region [%]<br>Region Type : Cytoplasm Region<br>Outer Border : -10 %<br>Inner Border : 55 % | Output Region : Cytoplasm Region |

| Calculate Intensity Properties (3) | Input | Method | Output |
|---|---|---|---|
| | **Channel :** MitoTracker Green<br>**Population :** Live cells<br>**Region :** Cytoplasm Region | **Method :** Standard Mean | Property Prefix : Intensity Cytoplasm Region MitoTracker Green |

| Calculate Intensity Properties (4) | Input | Method | Output |
|---|---|---|---|
| | **Channel :** TMRE<br>**Population :** Live cells<br>**Region :** Cytoplasm Region | **Method :** Standard Mean | Property Prefix : Intensity Cytoplasm Region TMRE |

| Calculate Intensity Properties (5) | Input | Method | Output |
|---|---|---|---|
| | **Channel :** MitoTracker Green<br>**Population :** Live cells<br>**Region :** Cell | **Method :** Standard Mean | Property Prefix : Intensity Cell MitoTracker Green |

| Calculate Intensity Properties (6) | Input | Method | Output |
|---|---|---|---|
| | **Channel :** TMRE<br>**Population :** Live cells<br>**Region :** Cell | **Method :** Standard Mean | Property Prefix : Intensity Cell TMRE |

| Calculate Properties | Input | Method | Output |
| --- | --- | --- | --- |
| | **Population :** Live cells | **Method :** By Formula<br>Formula : A/B<br>Variable A : Intensity Cytoplasm Region TMRE Mean<br>Variable B : Intensity Cytoplasm Region MitoTracker Green Mean | Output Property : Norm. Cytoplasmic TMRM |

| Define Results | Results |
| --- | --- |
| | **Method :** List of Outputs<br>**Population : Live cells**<br>Number of Objects<br>Intensity Cytoplasm Region MitoTracker Green Mean : Mean+StdDev<br>Intensity Cytoplasm Region TMRE Mean : Mean+StdDev<br>Intensity Cell MitoTracker Green Mean : Mean+StdDev<br>Intensity Cell TMRE Mean (2) : Mean+StdDev<br>Norm. Cytoplasmic TMRM : Mean+StdDev<br><br>**Object Results**<br>Population : True Nuclei : None<br>Population : Nuclei : None<br>Population : Live cells : ALL |

### References

1. Forny, P. *et al.* Integrated multi-omics reveals anaplerotic rewiring in methylmalonyl-CoA mutase deficiency. *Nat. Metab.* **5**, 80–95 (2023).
2. Luciani, A. *et al.* Impaired mitophagy links mitochondrial disease to epithelial stress in methylmalonyl-CoA mutase deficiency. *Nat. Commun.* **11**, 970 (2020).
3. Köpfer, F. *et al.* Effects of anserine on oxidative stress and on cell barrier integrity in methylmalonic aciduria. *Sci. Rep.* **15**, 32933 (2025).
4. Denley, M. C. S. *et al.* Mitochondrial dysfunction drives a neuronal exhaustion phenotype in methylmalonic aciduria. Preprint at https://doi.org/10.1101/2024.03.15.585183 (2024).
5. Anzalone, A. V. *et al.* Search-and-replace genome editing without double-strand breaks or donor DNA. *Nature* **576**, 149–157 (2019).
6. Chen, P. J. *et al.* Enhanced prime editing systems by manipulating cellular determinants of editing outcomes. *Cell* **184**, 5635-5652.e29 (2021).
7. Nelson, J. W. *et al.* Engineered pegRNAs improve prime editing efficiency. *Nat. Biotechnol.* **40**, 402–410 (2022).
8. Tálas, A. *et al.* BEAR reveals that increased fidelity variants can successfully reduce the mismatch tolerance of adenine but not cytosine base editors. *Nat. Commun.* **12**, 6353 (2021).
9. Zhang, Y. *et al.* Rapid Single-Step Induction of Functional Neurons from Human Pluripotent Stem Cells. *Neuron* **78**, 785–798 (2013).
10. Hockemeyer, D. *et al.* A Drug-Inducible System for Direct Reprogramming of Human Somatic Cells to Pluripotency. *Cell Stem Cell* **3**, 346–353 (2008).
11. Dull, T. *et al.* A Third-Generation Lentivirus Vector with a Conditional Packaging System. *J. Virol.* **72**, 8463–8471 (1998).
12. Buescher, J. M. *et al.* A roadmap for interpreting 13C metabolite labeling patterns from cells. *Curr. Opin. Biotechnol.* **34**, 189–201 (2015).
